# The long noncoding RNA *Peanut* (*Gm11454*) promotes neurogenesis and rod photoreceptor differentiation during postnatal retinal development

**DOI:** 10.64898/2025.12.02.691925

**Authors:** Jade Enright Hostetler, Fion Shiau, Xiaodong Zhang, Shiming Chen, Philip A. Ruzycki, Seth Blackshaw, Brian S. Clark

**Author notes:** Current Address: Department of Systems Biology, Columbia University Irving Medical Center, New York, NY 10032 and the New York Genome Center, New York, NY 10013. Corresponding Author: Brian Clark.

## Abstract

Long noncoding RNAs (lncRNAs) display pervasive expression and function in the developing nervous system. Temporal profiling of gene expression in the retina has demonstrated differential expression of lncRNAs throughout development; however, determinations of lncRNA function during retinal development remain limited. In this study, we identify numerous lncRNAs with dynamic temporal expression and characterize the function of the lncRNA *Gm11454,* which we have named *Peanut*. Using overexpression of *Peanut* in mice retinas, we determine that *Peanut* promotes rod photoreceptor fate and neurogenesis of retinal progenitor cells (RPCs) via inhibition of Notch signaling and by regulating expression of neighboring gene *Tox2*. A novel *Peanut* knockout mouse model demonstrates that *Peanut* is required for proper visual function and photoreceptor gene expression. Finally, we determined that *Peanut* is necessary for proper cell cycle progression and neurogenesis. Our results characterize the function of a novel lncRNA as a regulator of RPC neurogenesis and differentiation and support the importance of lncRNAs in the developing retina.

## 1. Introduction

The vertebrate retina is comprised of seven major cell types arranged across three nuclear layers that transform visual stimuli into electrical signals that are relayed to the brain. The seven cell types – six neuronal and one glial – arise from a single pool of retinal progenitor cells (RPCs; Turner and Cepko 1987; Turner et al.,1990; reviewed in Bassett and Wallace, 2012) that either proliferate to expand the progenitor population or undergo neurogenic and, later, gliogenic divisions for terminal cell fate differentiation (Centanin and Wittbrodt, 2014). The seven major cell classes of the mature retina are generated in a conserved overlapping temporal order during development; retinal ganglion cells, horizontal cells, cone photoreceptors and amacrine cells differentiate in an early, mostly embryonic wave in mice, followed by a postnatal wave where rod photoreceptors, bipolar cells, and Müller glia are generated (Stenkamp, 2015). Numerous studies have identified key transcription factors and epigenetic mechanisms that dictate retinal neurogenesis and cell fate (reviewed in Shiau et al., 2021; Raeisossadati et al., 2021); however, the nature of precise temporal control of gene regulatory networks (GRNs) and temporal cell fate specification remains incomplete. The simple organization and accessibility of the retina provide a valuable model for understanding the molecular dynamics underlying nervous system development.

Increasing evidence highlights the pervasive expression and function of long noncoding RNAs (lncRNAs) in nervous system development. LncRNAs are transcripts of over 200 nucleotides that lack significant protein coding potential (Statello et al., 2021). LncRNAs regulate various biological processes by acting as molecular scaffolds, regulating gene expression and cellular processes through a variety of mechanisms including chromatin organization, RNA-protein scaffolding, and miRNA sponging (reviewed in Clark and Blackshaw, 2014; Lin et al., 2014; Engreitz et al., 2016; Rani et al., 2016; Li et al., 2025). Additionally, lncRNAs have high tissue and cell type specificity compared to protein coding genes – on average, 50% of protein coding genes are expressed in individual cell types (Hezroni et al., 2019). Conversely, individual cell types express less than 5% of annotated lncRNAs (Hezroni et al., 2019). LncRNAs display enriched expression within the central nervous system, and numerous lncRNAs have been characterized during brain development (Hamazaki et al., 2015; Mercer et al., 2008; Sauvageau et al., 2013). During corticogenesis, differentially expressed lncRNAs are found to act as molecular “switch” genes, biasing progenitors to transition from proliferative divisions to neurogenic divisions – where at least one daughter cell exits the cell cycle and differentiates as a neuron (Aprea et al., 2013). We hypothesize that lncRNAs play a similar role in the retina, regulating proliferative to neurogenic transitions of progenitors and impacting cell fate decisions.

Temporal profiling of gene expression in the retina has demonstrated differential expression of lncRNAs throughout development and aging (Kong et al., 2023). Recent studies determined that 4,500 lncRNAs are expressed during mouse retinal development, often with spatial and temporal specificity (Yu et al., 2023). Additionally, a population of photoreceptor-specific lncRNAs has been identified and shown to shift in expression dynamics during development (Zelinger et al., 2017). Despite their prevalent expression, determinations of lncRNA function during retinal development remain limited: *Vax2os* and *Tug1* are necessary for the cell cycle progression and formation of photoreceptor precursors, respectively (Meola et al., 2012; Young et al., 2005). *RNCR2 (Miat)* regulates cell cycle exit, amacrine cell and Müller glia differentiation, whereas *Six3os1* regulates the activity of the transcription factor Six3 during retinal cell fate specification through interactions with Eya proteins, which function as Six3 co-activators (Rapicavoli et al., 2010, 2011). Beyond previous studies assessing the function of *Six3os1*, the mechanisms by which lncRNAs function in the retina remain limited.

To prioritize candidate lncRNA regulators of retinal development, we utilized previously published RNA expression profiling to identify dynamically expressed lncRNAs with expression patterns consistent with a neurogenic switch of RPCs and the temporal specification of retinal cell fates, validating the spatio-temporal expression pattern of a subset of temporally expressed lncRNAs. We further assessed the expression and function of an uncharacterized lncRNA *Gm11454*, which we named *<u>P</u>hotor<u>E</u>ceptor <u>A</u>nd <u>N</u>eurogenesis <u>U</u>pregulating <u>T</u>ranscript (Peanut).* Using overexpression and knockout mouse models, we demonstrate that *Peanut* regulates cell cycle progression and differentiation of photoreceptors, identifying potential mechanisms of *Peanut* function.

## 2. Results

### 2.1 LncRNAs are differentially expressed in the developing retina and display spatiotemporal and cell type specific expression patterns

We first sought to investigate broad patterns of lncRNA expression in the developing retina. In previous studies, we utilized a developmental time-series profiling temporal transcript expression patterns to identify novel regulators of retinal cell fate specification (Clark et al., 2019). We hypothesized that identification of lncRNAs with temporally dynamic expression profiles would likewise uncover novel lncRNA regulators of retinal development. To achieve this, we analyzed a temporal series of bulk RNA-seq data of *Chx10*-EGFP/cre-ALPP (Rowan and Cepko, 2004) fluorescence-activated cell sorted (FACS) RPCs and post-mitotic cells (PMCs) at (E)mbryonic day 14 (early development) and (P)ostnatal day 2 (late development) (Stein-O’Brien et al., 2019; Zibetti et al., 2019; Figure 1A). We hypothesized that lncRNAs enriched within RPCs function to control RPC proliferative potential. We additionally postulate that RPC-enriched lncRNAs with temporally dynamic expression across early and late progenitor datasets have the potential to regulate the temporal specification of retinal cell fates. Our transcriptomic analyses identified numerous dynamically expressed lncRNAs (Figure S1) whose retinal expression patterns were confirmed using a time-series of chromogenic *in situ* hybridization (Table S1). We characterized the expression of individual lncRNAs that displayed enrichment in early (e.g., *Gm29266*, *Gm16551*) or late (e.g., *Sox8os, 2610035D17Rik*) development periods (Figure 1B, C). Notably, identified lncRNAs included previously characterized lncRNAs like *Miat* (Rapicavoli et al., 2010, Figure S1), uncharacterized lncRNAs transcribed from the opposite strand of transcription factors (*Prox1os, Nfia-os, Sox8os;* Figures 1B, C and S1), and uncharacterized intergenic lncRNAs (*Gm29266, Gm16551,* Figures 1B, C).

**Figure 1.**
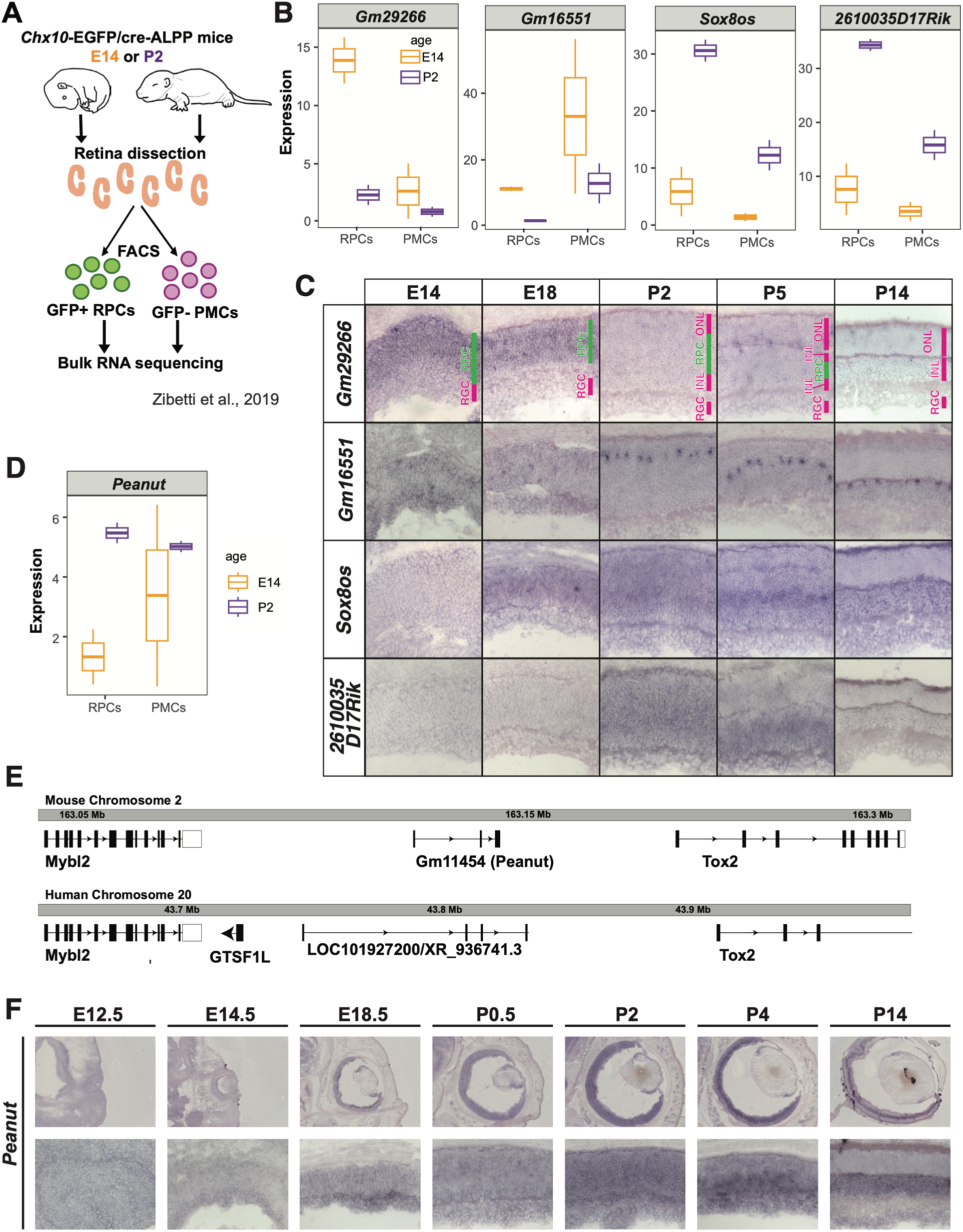
LncRNAs are specifically and dynamically expressed in the developing retina. A) Experimental design for sorting and sequencing of developmental time-series RNA-seq, as used in Zibetti et al., 2019. B) Average expression of individual lncRNAs in RPCs and PMCs at E14 (orange) and P2 (purple) in RNA-seq experiments. C) Chromogenic *in situ* hybridization time series profiling spatiotemporal expression patterns of individual lncRNAs. D) Expression of *Peanut,* detected by RNA-seq analysis in RPCs and PMCs at E14 (orange) and P2 (purple). E) The genomic locus of *Peanut* (mouse) and its syntenic human transcript, containing conserved sequence homology with the *Peanut* transcript. F) ISH across development of *Peanut*.

We additionally assessed cellular specificity of lncRNAs in comparison to protein coding genes across our previously published developing retina single cell (sc) RNA sequencing (RNA-seq) datasets (Clark et al., 2019). Consistent with previous reports detailing the cellular specificity of lncRNA transcripts (Hezroni et al., 2019), a high proportion of lncRNA transcripts were expressed in individual major retinal cell types. Conversely, a high proportion of protein-coding transcripts were ubiquitously expressed across cell types (Figure S2). lncRNA cell type specificity was further analyzed across scRNA-seq datasets of mature retinal ganglion cell and amacrine cell subtypes (Tran et al., 2019; Yan et al., 2020). Similarly, lncRNA transcript expression was enriched within individual cell subtypes, while protein coding transcripts were more often expressed across cell subtypes (Figure S2). Notably, scRNA-seq analysis fails to detect non-polyadenylated lncRNAs, as well as lncRNAs with expression below dropout detection thresholds; however, our analyses at the cell type or cellular subtype level demonstrate that lncRNAs were expressed at similar levels to protein coding gene transcripts when using a capture threshold of at least 5% of cells in each cell type (Figure S3). The high cellular specificity of lncRNAs in the retina supports the hypothesis that lncRNAs exhibit cell type specificity with the potential to influence specification and differentiation of individual cell fates.

Within our examinations of temporally dynamic gene expression across retinal development, we identified the uncharacterized lncRNA *Gm11454,* which we have named *Photoreceptor and neurogenesis upregulating transcript (Peanut),* given our findings described below. Our temporal RNA-seq analysis in the developing retina demonstrated enriched expression of *Peanut* in early PMCs compared to early RPCs, and significant enrichment of *Peanut* expression in P2 RPCs compared to E14 RPCs (Figure 1D). *Peanut* is transcribed as an intergenic lncRNA from mouse chromosome 2 between the genes *Mybl2* and *Tox2* (Figure 1E). We observe low sequence conservation of *Peanut* (128 bp with 82.9% sequence) within the locus of a syntenic human transcript (LOC101927200/XR_936741.3; Figure 1E). lncRNAs evolve rapidly and lack primary sequence conservation across species; however, syntenic transcripts are often observed, suggesting functional conservation of the RNA transcript in a sequence-independent manner (Diederichs, 2014). *In situ* hybridization targeting of the *Peanut* transcript in the developing retina identified *Peanut* expression in early retinal development, notably in the presumptive retinal ganglion cell layer and inner nuclear layer, consistent with the RNA-seq analysis results. *Peanut* expression in the retina increased at P2, exhibiting broad expression in both the neuroblast layer and within post-mitotic cells. By P14, *Peanut* expression is restricted to the inner nuclear and RGC layers (Figure 1F).

### 2.2 Overexpression of Peanut alters retinal cell fate specification and RPC proliferation

We next utilized overexpression studies to test the sufficiency of *Peanut* for the regulation of retinal cell fate specification and RPC cell cycle exit. *In vivo* electroporation at P0 followed by immunohistochemistry at P14 for cell type specific markers (Figure 2A, Table S2) revealed that *Peanut* overexpression biased RPCs to generate rod photoreceptors at the expense of bipolar cells, amacrine cells, and Müller glia compared to empty vector control electroporations (Figure 2B, C).

**Figure 2.**
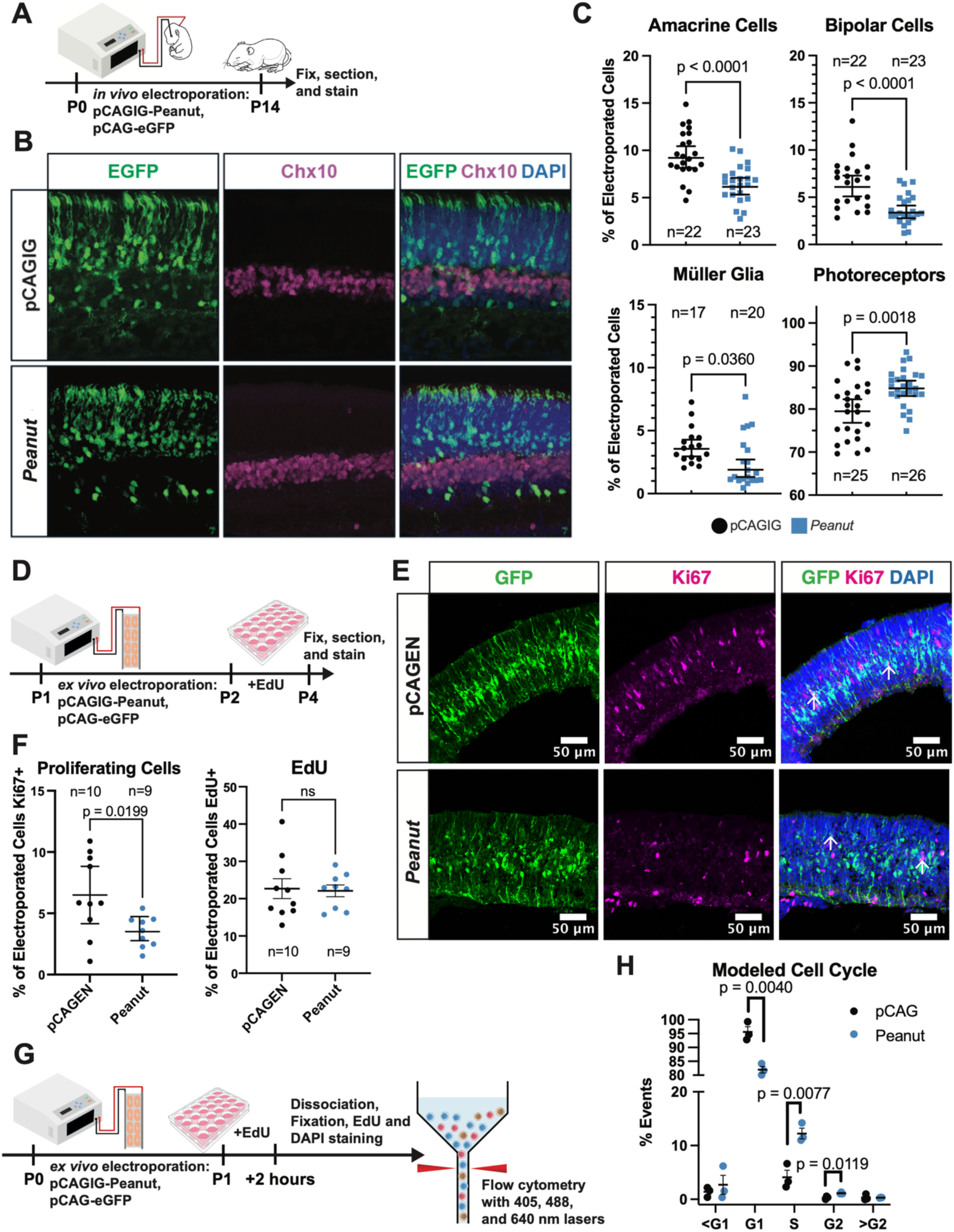
Overexpression of Peanut promotes photoreceptor fate and neurogenesis. A) P0 mice were *in vivo* electroporated with pCAG-Peanut or empty pCAGIG and co-electroporated with pCAG-emGFP. Eyes were fixed at P14, processed for sectioning and immunohistochemistry staining for cell fate markers. B) Control and *Peanut*-overexpressing retinas stained for GFP (electroporation marker), Chx10 (Vsx2, bipolar cell marker), and DAPI (nuclei). C) Quantification of cell fate markers in *Peanut* and control pCAGIG electroporations. The number of cells positive for a cell fate marker was divided by the total number of electroporated cells to calculate percent cell fate and normalize for electroporation efficiency. Counts were performed in triplicate images per retina with n’s representing individual retinas. Mean ± SEM displayed, statistical significance determined by student’s t-test. D) P1 mice retinas were explanted and electroporated with pCAGEN-*Peanut* or empty pCAGEN and pCAG-emGFP. After 24 hours, a 48-hour EdU pulse was performed. Retinal explants were then fixed, sectioned, and stained. E) Control and *Peanut*-overexpressing retinas stained for GFP (electroporation marker), Ki67 (proliferative marker), and DAPI. F) Quantification of the percent of electroporated cells that were Ki67+ and EdU+. Counts were performed over triplicate images per retina. Mean ± SEM displayed, statistical significance determined by student’s t-test, ns = not significant. G) P0 mice retinas were explanted and electroporated. After 24 hours, a 2-hour EdU pulse was performed followed by dissociation, fixation, and staining for EdU and DAPI. Cells were then used for flow cytometry analysis of electroporated cells (GFP+) for EdU accumulation and DNA content. H) Percent of cells in each cell cycle phase as determined by cell cycle modeling of control and Peanut-overexpressing cells using FlowJo cell cycle tool. <G1 and >G2 represent cells outside of the G1 and G2 peaks with less or more DNA content, respectively. Mean ± SEM displayed, statistical significance determined by student’s t-test, ns = not significant.

We also assessed the impact of *Peanut* overexpression on RPC cell cycle exit using EdU incorporation assays. One day after P0 *ex vivo* electroporation, a 24-hour or 48-hour pulse of EdU was administered to track cycling cells (progressing through S-phase) and the extent to which *Peanut* regulates terminal divisions of RPCs (Figure 2D). We assessed the proportions of electroporated cells that 1) passed through S-phase (incorporated EdU), and 2) remained proliferative (MKi67+) after *Peanut* electroporation. *Peanut* overexpression did not affect EdU accumulation after 24-hours or 48-hours, indicating there is not a significant difference in the number of RPCs proceeding through the cell cycle (Figure 2F, S4). Ki67 staining also was not affected when retinas were fixed 48 hours after *Peanut* overexpression (Figure S4). However, when retinas were fixed 72 hours after electroporation, *Peanut-*overexpressing cells were significantly less likely to co-stain for Ki67 (Figure 2E, F), suggesting that *Peanut* overexpression biases RPCs to exit the cell cycle.

To further investigate the role of *Peanut* in cell cycle regulation, flow cytometry analysis for DNA content was performed on retinal explants overexpressing *Peanut* or control empty vector. 24 hours after electroporation, a 2-hour EdU pulse was performed to mark cells in S-phase, followed by dissociation, fixation, and staining for EdU and DAPI (Figure 2G). Flow cytometry and cell cycle modeling of electroporated cells based on DAPI signal demonstrated that *Peanut-*overexpressing cells were less often in the G1 phase, and more often in the S and G2 phases of the cell cycle (Figure 2H, S5). *Peanut-*overexpressing cells were also more likely to be EdU-positive, consistent with the increase in the S phase population (Figure S4, S5).

This finding is in opposition to our observation of unchanged EdU accumulation over a 24-hour or 48-hour pulse (Figure 2F), which may be explained by the heightened sensitivity of flow cytometry detection and the pulse length. EdU accumulation over 2 hours (increased after *Peanut* overexpression) represents the number of cells actively in the S-phase, while EdU accumulation over 24 or 48 hours (unchanged after *Peanut* overexpression) represents the number of cells which passed through S phase (actively cycling) during the EdU pulse. So, while *Peanut* overexpression lengthens S-phase, it does not affect the number of cells actively cycling. Additionally, a lengthened G2 phase biases progenitor cells towards more terminal, neurogenic divisions (Calegari et al., 2005), consistent with our observation of fewer proliferative cells following *Peanut* overexpression (Figure 2E, F). Thus, *Peanut* overexpression does not directly cause cell cycle exit but influences cell cycle phase progression to promote neurogenic divisions over self-renewing divisions. Overall, we conclude that *Peanut* overexpression biases RPCs towards neurogenic divisions in the early postnatal retina to promote photoreceptor fate.

### 2.3 Peanut overexpression inhibits Notch signaling and Tox2 expression

We next sought to investigate the molecular mechanism by which *Peanut* overexpression promotes neurogenesis and photoreceptor fate in RPCs. Identification of the subcellular localization of lncRNAs assists in determining putative lncRNA functions. For example, many nuclear-enriched lncRNAs participate in the regulation of chromatin structure and/or gene expression (Lin et al., 2014; Engreitz et al., 2016; Rani et al., 2016; Li et al., 2025). Therefore, to assess the putative functions of *Peanut* we utilized multiple orthogonal approaches to identify and confirm *Peanut* subcellular localization. Using quantitative, reverse transcription PCR (qRT-PCR) on isolated subcellular fractionations of *Peanut-*transfected NIH3T3 cells, we determined that *Peanut* was enriched in the salt-extracted (loosely bound to chromatin) and chromatin-bound fractions (Figure 3A, S6; Table S3). The specificity of subcellular fractionations was highlighted by nuclear (specifically salt-extracted fraction) enrichment of *Malat1*, a lncRNA shown to be enriched in a nuclear-insoluble fraction (Chen et al., 2017; Figure S6). We further confirmed nuclear enrichment of *Peanut* using an MS2 aptamer-based approach in HEK293 cells (Bertrand et al 1998, Fusco et al 2003). MS2 stem loops were cloned into the 3’ end of the transcript sequences of *Peanut* or *tdTomato* and co-transfected with the MS2 coat protein conjugated to GFP for MS2-containing transcript localization (Figure S6). The specificity of the MS2 approach was first determined using *tdTomato* transcript as a control. *tdTomato* MS2-GFP fluorescence strongly overlapped with red (tdTomato) fluorescence, both in the cytoplasm and nucleus (Figure S6). However, we observed strong localization of GFP to the nucleus, indicating a nuclear enrichment of *Peanut* (Figure S6). Combined, our results identify that *Peanut* is enriched in the nucleus and interacts with chromatin, positioning *Peanut* to regulate transcription, chromatin structure, or recruitment of epigenetic modifiers to DNA. Based on both spatiotemporal expression and nuclear enrichment of *Peanut*, we hypothesize that *Peanut* influences gene expression to regulate cell cycle exit or cell fate specification during retinal development.

**Figure 3.**
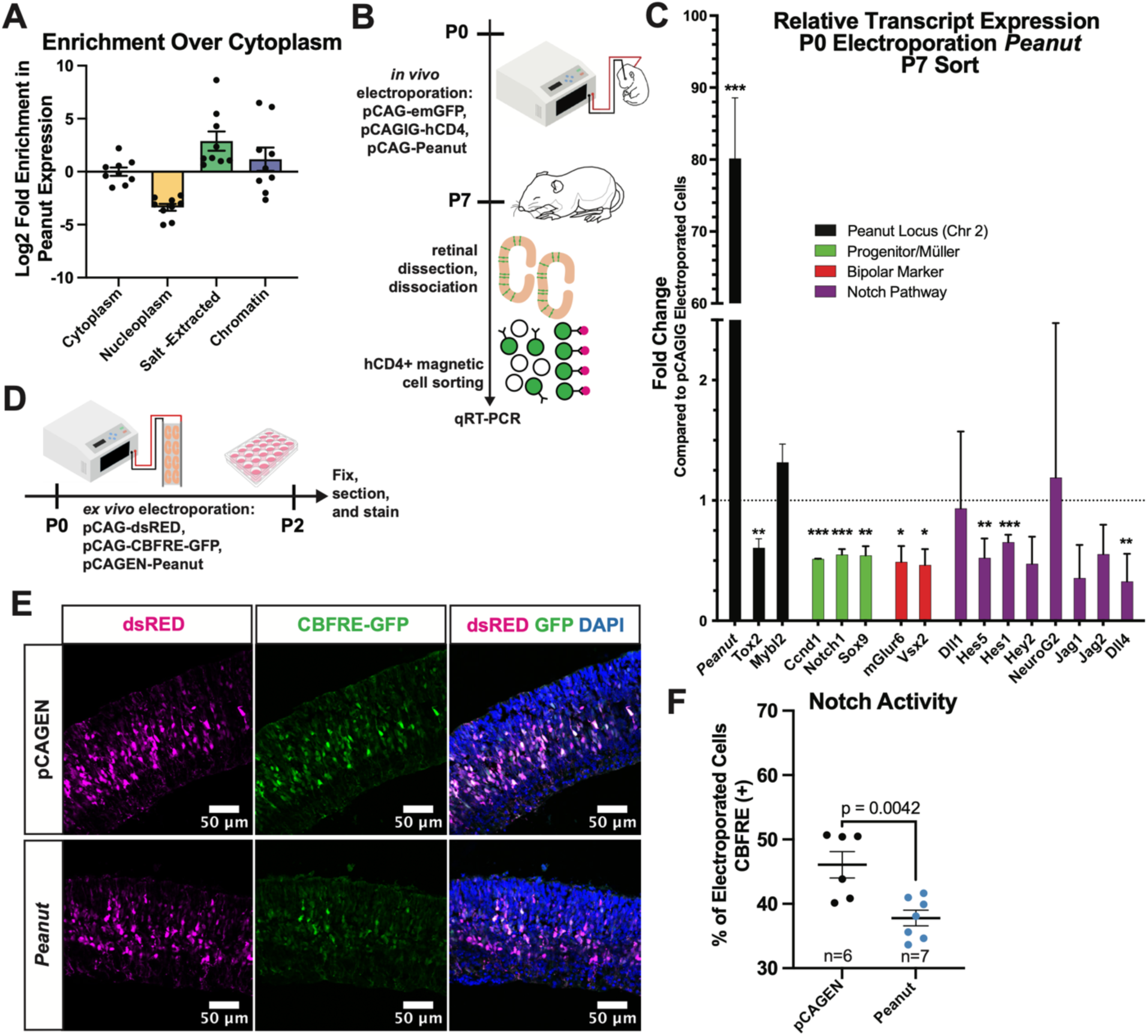
Peanut overexpression inhibits Notch signaling. A) Subcellular fractionation of NIH3T3 cells followed by qRT-PCR of *Peanut.* After cytoplasmic removal generating the Cytoplasm fraction, nuclei were lysed and supernatant removed as the Nucleoplasm fraction. The pellet was then separated into Salt-Extracted (loosely associated with chromatin) and Chromatin fractions. Shown is enrichment of each fraction over the cytoplasmic fraction. B) pCAG-emGFP and empty pCAGIG control or pCAG-Peanut were co-electroporated in P0 mice with pCAGIG-hCD4. At P7, retinas were dissected and dissociated. hCD4+ magnetic cell sorting was performed to enrich for electroporated cells, which were used for cDNA synthesis and qRT-PCR. C) Fold-change of transcript expression in *Peanut-*overexpressing retinas compared to control. D) P0 retinal explants were electroporated with pCAGEN-*Peanut* or control empty pCAGEN, pCAG-dsRED, and pCAG-CBFRE (GFP) and fixed at P2, followed by sectioning and staining. E) Retinas were stained for dsRED (electroporation marker), GFP (CBFRE to report Notch pathway activity), and DAPI (nuclei). F) Quantification of percent of electroporated cells positive for Notch activity as indicated by CBFRE. Counts were performed across triplicate images per retina. Mean ± SEM displayed, statistics determined by student’s t-test. N’s represent results from individual retinal explants.

To understand the gene expression changes that occur following *Peanut* overexpression, we profiled gene expression changes in P0 *Peanut* electroporated cells by qRT-PCR at P7. To enrich for electroporated cells, we utilized magnetic-activated cell sorting by co-electroporating a pCAGIG-human CD4 plasmid (hCD4). CD4 is not expressed in the developing retina, and surface membrane expression of hCD4 enables pulldown of electroporated cells using anti-hCD4 antibody conjugated beads (Figure 3B). We assessed *Peanut*-induced changes in gene expression using a qRT-PCR gene panel for cell type-specific markers and components of the Notch signaling pathway – a pathway known to influence both retinal neurogenesis and cell fate specification (Table S3). Markers of progenitor cells (*Ccnd1, Notch1, Sox9*) and bipolar cell fate (*mGlur6, Vsx2*) showed decreased expression levels, consistent with the observed decrease in proliferation and decreased bipolar cell fate at P14 (Figure 2, 3C). Interestingly, *Peanut* overexpression led to decreased levels of several Notch signaling pathway transcripts (Figure 3C). Notch signaling is required for continued proliferation of RPCs and must be downregulated for cell cycle exit and neurogenesis (Jadhav et al., 2006a; Mills and Goldman, 2017). Notably, Notch signaling also inhibits photoreceptor formation during retinal development (Jadhav et al., 2006b; Yaron et al., 2006). This finding presented a potential mechanism of action for *Peanut* whereby *Peanut* inhibits Notch signaling, leading to increased neurogenesis and photoreceptor differentiation. To confirm that *Peanut* leads to a decrease in Notch pathway activation, we co-electroporated the CBF1 Responsive Element (CBFRE) Notch reporter with *Peanut* (Figure 3D). The CBFRE reporter construct contains four CBF1 Notch-responsive Rbpj binding sites upstream of GFP, providing a functional readout of pathway activity (Nowotschin et al., 2013). We observed that overexpression of *Peanut* led to a decrease in Notch-responsive, GFP+ cells, demonstrating that *Peanut* inhibits Notch signaling (Figure 3DE, F).

In our qRT-PCR gene panel, we also assessed the expression of transcripts within the *Peanut* gene locus, as lncRNAs frequently regulate the gene locus from which they are transcribed and participate in the same regulatory pathways (Gil and Ulitsky, 2019). *Peanut* overexpression led to a decrease in *Tox2* while *Mybl2* expression remained unaffected (Figure 3C). Tox2 is a transcription factor whose function in the developing retina remains uncharacterized. *In situ* hybridization for *Tox2* expression within the developing retina indicated broad expression in the early developing retina across both RPCs and PMCs. However, beginning at P2, *Tox2* expression progressively restricts to the inner nuclear layer and ganglion cell layer (Figure 4A, B).

**Figure 4.**
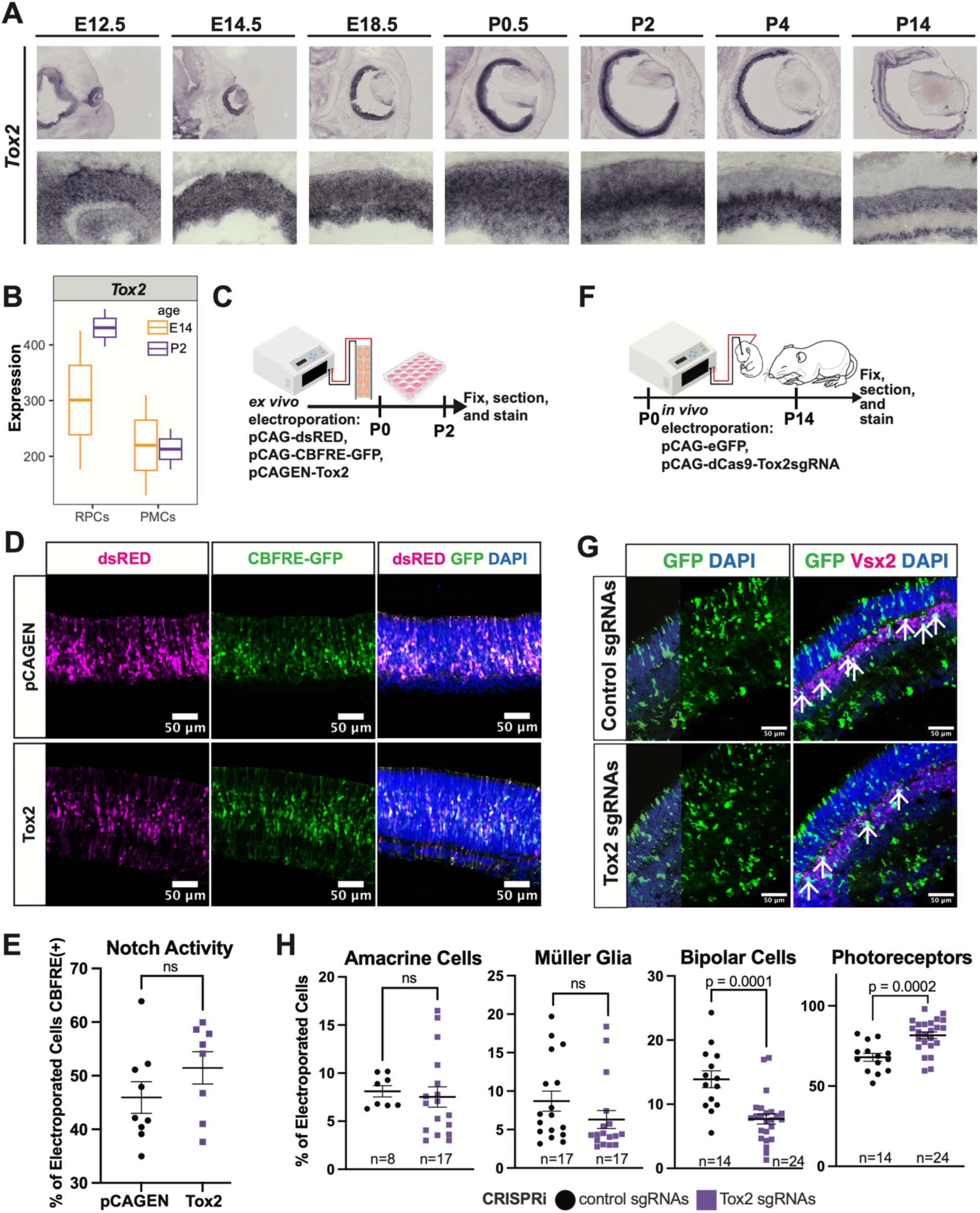
Tox2 is required for proper cell fate proportions but does not regulate Notch signaling. A) *In situ hybridization* across development of *Tox2.* B) Expression of *Tox2*, detected by RNA-seq analysis in RPCs and PMCs at E14 (orange) and P2 (purple). C) P0 retinal explants were electroporated with pCAGEN-*Tox2* or empty pcAGEN, pCAG-dsRED, and pCAG-CBFRE (GFP), and fixed at P2, followed by sectioning and staining. D) Retinas were stained for dsRED (electroporation marker), CBFRE (GFP, Notch activity), and DAPI (nuclei). E) Quantification of percent of electroporated cells positive for CBFRE. Counts were performed across triplicate images per retina. Mean ± SEM displayed, statistics determined by student’s t-test with n’s representing individual retinas. F) P0 mice were electroporated *in vivo* with pCAG-eGFP and pCAG-dCas9 with *Tox2* sgRNAs or pCAG-dCas9 with control sgRNAs. At P14, eyes were fixed, sectioned, and stained. G) Staining of *Tox2*– and control-CRISPRi retinas with emGFP (electroporation marker), Chx10 (Vsx2, bipolar cell fate marker), and DAPI (nuclei). Arrows indicate co-stained cells (GFP+ Vsx2+) H) Quantification of cell fate marker staining in *Tox2-* and control-CRISPRi electroporations. Counts were performed across triplicate images per retina with n’s representing retinas from individual animals. Mean ± SEM displayed, statistics determined by student’s t-test.

We hypothesized that Tox2 promotes Notch activity, and therefore *Peanut* functions via *Tox2* expression regulation to attenuate Notch signaling. We overexpressed *Tox2* and assessed Notch pathway activation using the CBFRE-GFP reporter (Figure 4C). Tox2 overexpression failed to increase the autonomous activation of the CBFRE reporter compared to control electroporated cells (Figure 4D, E), implying that Tox2 does not directly regulate Notch pathway activation. To further investigate the role of Tox2 in the developing retina, *Tox2* expression was inhibited utilizing *in vivo* electroporation of the CRISPRi system (Gilbert et al., 2013, Figure 4F). A guide RNA (gRNA) targeting the first exon of *Tox2* driven by the *U6* promoter (pMU6-Tox2 CRISPRi) was co-electroporated with a construct encoding the dCas9-KRAB fusion protein (pCAG-dCas9-BFP-KRAB). Loss of *Tox2* expression in the developing retina increased the number of rod photoreceptor cells and correspondingly decreased the number of bipolar cells (Figure 4G, H). Additionally, there was a trending decrease in amacrine cells and Müller glia in *Tox2-CRISPRi* electroporated cells (Figure 4H). This phenotype mimics *Peanut* overexpression, suggesting that *Peanut* promotes photoreceptor specification of postnatal RPCs by reducing *Tox2* expression and/or function.

### 2.4 Loss of Peanut impacts photoreceptor function

To investigate the necessity of *Peanut* during retinal development, we next generated a loss-of-function mouse model. A∼25kb deletion of the entire *Peanut* locus was generated using the CRISPR/Cas9 system, with guide RNAs flanking the *Peanut* 5’-most and 3’-most exons (*Peanut^−/−^*; Figure 5A). Knockout animals are viable and healthy, with *Peanut* mutants obtained in expected ratios (Table S4). Given that *Peanut* overexpression increased photoreceptor fate specification, we hypothesized that *Peanut^−/−^* mice would display decreased numbers of rod photoreceptors and exhibit decreased visual function. To assess visual function, we performed electroretinography (ERG) on 8-week-old wildtype (WT) control, *Peanut* heterozygote (*Peanut^+/−^)*, and *Peanut^−/−^* mice. Both scotopic A-wave and scotopic B-wave were slightly but significantly decreased in *Peanut^−/−^* mice compared to both *Peanut^+/−^* and WT mice at the brightest scotopic intensities (Figure 5B). These results suggest a decrease in rod photoreceptor function, consistent with our predictions. However, *Peanut^−/−^* retinas did not display significant differences in retinal gross morphology or retinal thickness as assessed by H&E staining at P21, 8 weeks, and 1 year of age (Figure 5C, S7). To examine cell type proportions, we stained *Peanut^−/−^, Peanut^+/−^*, and WT retinas at P14, P22, 8 weeks, and 1 year by immunofluorescence with established cell type markers (Figure 5D, S8; Table S3). We did not observe significant differences in the proportion of photoreceptor cells or any other cell fate following *Peanut* loss of function (Figure 5D, S8).

**Figure 5.**
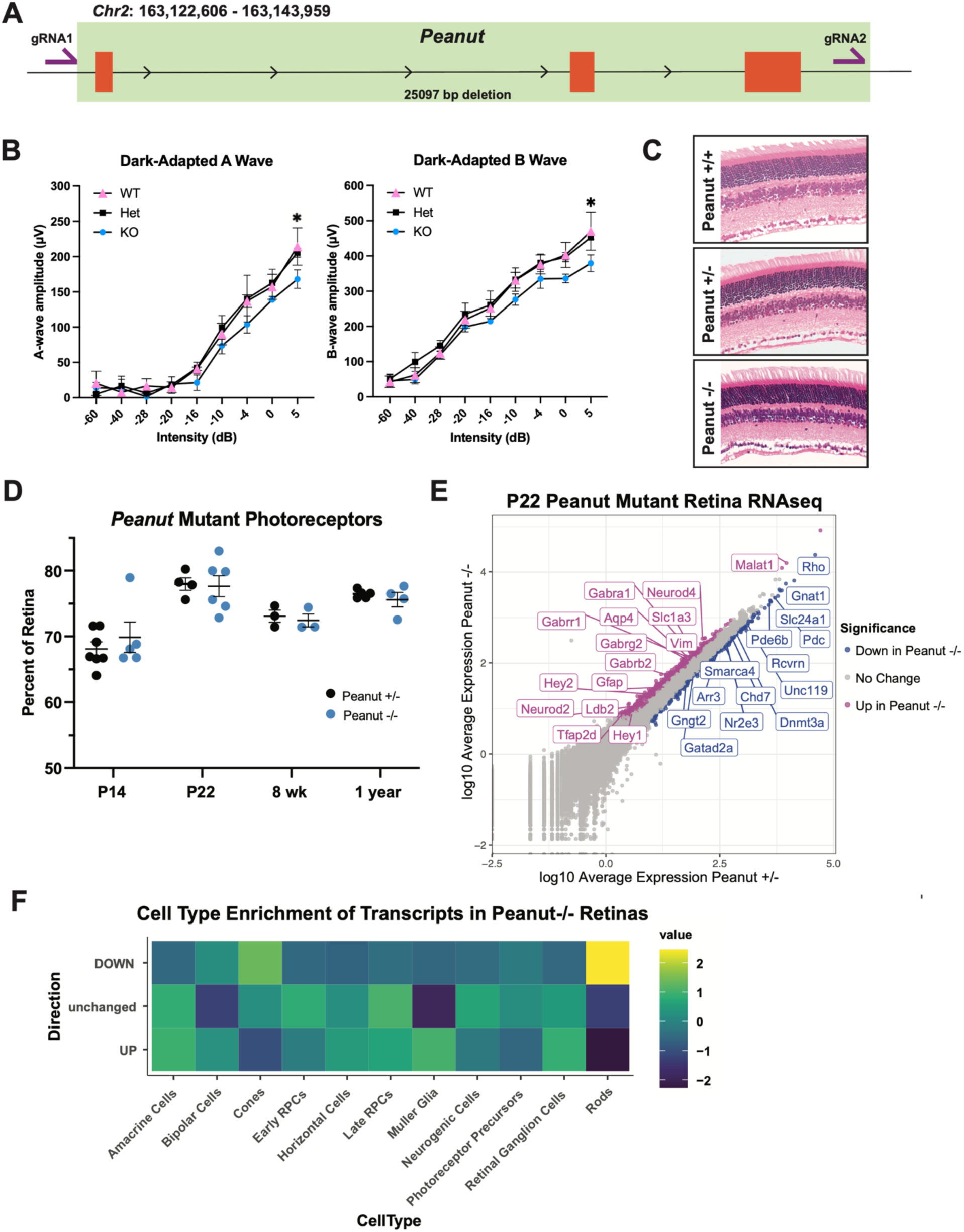
Peanut is required for proper photoreceptor function and gene expression. A) The *Peanut* locus was removed using CRISPR/Cas9. Green shaded region indicates the genomic deletion with *Peanut* exonic sequences indicated in orange. B) Scotopic A-and B-wave ERG readings of Peanut WT (WT, n=4), Peanut^+/−^ (Het, n=3), and Peanut^−/−^(KO, n=3) mice at 8 weeks old. C) H&E staining of *Peanut* WT, *Peanut^+/−^, and Peanut^−/−^*retinas at 8 weeks. D) Quantification of proportion of photoreceptors in *Peanut^−/−^ and Peanut^+/−^* retinas: percent of retina positive for photoreceptor cell fate markers within a set area. Counts performed across triplicate images per retina. Mean ± SEM displayed, statistics (no significant differences observed) determined by student’s t-test. E) Transcript expression in P22 *Peanut^−/−^ and Peanut^+/−^* retinas, colored by differential expression significance (FDR < 0.05) and direction (magenta=up in *Peanut^−/−^*; blue = down in *Peanut^−/−^)*. F) Heatmap showing the relative expression enrichment downregulated, unchanged, and upregulated transcripts from P22 *Peanut^−/−^* retinas within individual cell types across the scRNA-seq dataset of the developing mouse retina (Clark, et al., 2019). Cell type enrichment was determined using differentially expressed transcripts as input into the aggregate_gene_expression function in Monocle3.

To further understand the cause of visual function deficit in *Peanut^−/−^* retinas despite the lack of morphological or cellular deficits, we performed bulk RNA-seq on *Peanut^−/−^ and Peanut^+/−^* retinas at P0 and P22, ages corresponding to the onset of high levels of *Peanut* expression (P0) or within a differentiated retina (P22). Consistent with the low levels of *Peanut* expression within the embryonic retina, we observed very few transcriptional differences between *Peanut^−/−^ and Peanut^+/−^* control retinas at P0, with only 9 downregulated genes and 8 upregulated genes (Figure S9, Table S5). However, at P21, *Peanut^−/−^* retinas displayed robust changes in gene expression compared to *Peanut^+/−^* controls, with 205 downregulated genes and 1362 upregulated genes (Figure 5E, S9, Table S6). Gene ontology (GO) analysis revealed that genes downregulated in *Peanut^−/−^* retinas were enriched in pathways including visual perception, detection of light, chromatin organization, and mRNA and DNA metabolic processes. The latter pathways may be indicative of a more direct function of *Peanut* in gene expression regulation. Most notably, however, downregulated transcripts in *Peanut^−/−^*retinas were enriched for photoreceptor transcripts, including *Gnat1* and *Recoverin* (Figure 5E). Differentially expressed genes in our dataset were mapped as gene modules to individual cell types using published scRNA-seq data of the developing retina (Clark et al., 2019). We confirmed that downregulated transcripts in *Peanut^−/−^* retinas were highly enriched for rod photoreceptor cell expression (Figure 5F). Therefore, although loss of *Peanut* does not impact the specification of photoreceptors, absence of the lncRNA leads to decreased photoreceptor gene expression at P22 and functional deficits in rod photoreceptors in 8-week-old mice.

Conversely, genes unchanged and upregulated in *Peanut^−/−^*retinas did not display discrete expression enrichment in any singular cell type (Figure 5F). GO pathway analysis identified that upregulated genes were enriched in synaptic signaling and organization, and cell adhesion and migration. Specifically, several genes involved in GABA and G-protein signaling pathways were upregulated in Peanut−/− retinas. Upregulated transcripts also included cell type markers for late born cell fates other than rods, such as *Slc1a3* (aka *Glast1*, Müller glia), and *Neurod4* (amacrine and bipolar cells) (Figure 5E). Overall, these results may be indicative that a loss of *Peanut* causes a developmental delay or failure to establish mature photoreceptor GRNs.

Additionally, some Notch effector genes/genes in pathways promoting Notch signaling, including *Hey1* and *Hey2* (Iso et al., 2003; Sahu et al., 2021) were upregulated in *Peanut^−/−^* retinas. However, expression of many canonical Notch genes (e.g., *Notch1*) remained unaffected. To examine the effect of *Peanut* loss on Notch activity, we electroporated P0 *Peanut^−/−^ and Peanut^+/−^* retinas with CBFRE. We failed to observe significant differences in the number of CBFRE reporter positive cells in P2 *Peanut^−/−^* retinas (Figure S10). Therefore, we conclude that *Peanut* is not necessary for global regulation of Notch signaling but is sufficient to attenuate Notch activity autonomously when *Peanut* is overexpressed (Figure 3).

### 2.5 Peanut loss of function alters RPC cell cycle

Given the impact of *Peanut* overexpression on cell cycle progression and proliferation, we investigated whether *Peanut* loss also alters RPC cell cycle progression. We performed flow cytometry analysis following a 2-hour EdU pulse in *Peanut^+/−^* and *Peanut^−/−^* retinas with EdU and DAPI staining, such that cells could be sorted based on DNA content (Figure 6A). Modeling of cell cycle stages showed a significant decrease in both G1 and G2 populations in *Peanut^−/−^* retinas and no change in S phase populations within the EdU+ population (Figure 6B, S10). We confirmed alterations in the proportion of *Peanut^−/−^* RPCs in specific cell cycle phases through assessments of pH3 staining. While we did not observe a significant change in the number of mitotic cells between *Peanut^+/−^* and *Peanut^−/−^*, punctate staining of pH3 in non-apical cells – a characteristic of G2 phase (Hendzel et al., 1997; López-Sánchez et al., 2005) – was quantified (Figure 6C, D). These pH3-stained G2 cells were significantly decreased in *Peanut^−/−^* retinas, consistent with the decreased propensity of RPCs to be detected in the G2 phase observed in flow cytometry experiments and supporting the G2 phase shortening in *Peanut*^−/−^ retinas.

**Figure 6.**
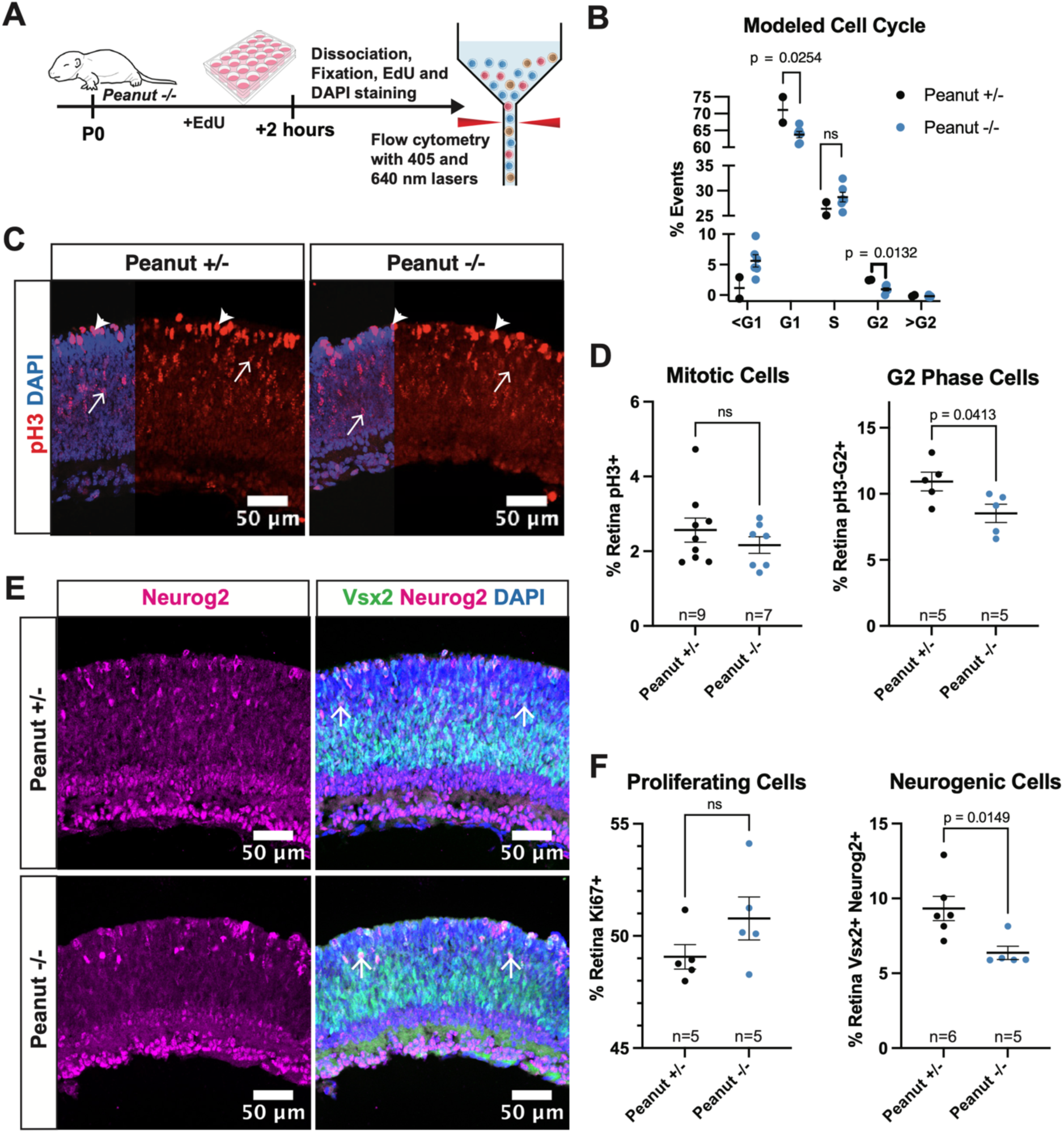
Peanut is required for proper cell cycle progression and neurogenesis. A) P0 *Peanut^+/−^*and *Peanut^−/−^* mice retinas were explanted, incubated with EdU for 2 hours, dissociated and fixed for staining. Cells were then used for flow cytometry analysis for EdU accumulation and DNA content. B) Percent of cells in each cell cycle phase as determined by cell cycle modeling using FlowJo cell cycle tool. <G1 and >G2 represent cells outside of the G1 and G2 peaks with less or more DNA content, respectively. Mean ± SEM displayed, statistical significance determined by student’s t-test. C) pH3 staining of *Peanut^+/−^* and *Peanut^−/−^* retinas showing mitotic cells (filled in arrowheads) and G2 cells (Open arrows). D) Quantification of pH3-stained mitotic and G2 cells. E) Neurog2 and Chx10/Vsx2 staining of *Peanut^+/−^* and *Peanut^−/−^* retinas. Co-staining examples shown with arrows. F) Quantification of proliferative cells as marked by Ki67 staining and neurogenic cells as marked by Ng2 and Vsx2 co-staining. Counts in D and F were performed as percent positive cells out of total retinal cells in set area, counts performed across triplicate images per retina. Mean ± SEM displayed, statistics determined by student’s t-test.

Shortening of either G1 or G2 phases biases progenitors to adopt self-renewing, proliferative divisions (Calegari and Huttner, 2003; Calegari et al., 2005; Peco et al., 2012). The decrease in RPCs in the G1 and G2 phases suggests that *Peanut^−/−^*retinas have an increase in self-renewing divisions and a decrease in neurogenic divisions, a complementary phenotype to those observed in *Peanut-*overexpressing retinas. To confirm a decrease in neurogenic divisions we co-stained *Peanut* mutant retinas with Neurog2 and Vsx2. Neurog2 is expressed in neurogenic RPCs (Hufnagel et al., 2010; Kowalchuk et al., 2018), with protein expression maintained in RGCs and amacrine cells. Co-labeling of Neurog2 with Vsx2 during development marks neurogenic RPCs. P0 *Peanut^−/−^* retinas displayed significantly decreased neurogenic RPCs as marked by co-staining of Neurog2 and Vsx2 (Figure 6E, F) as well as trending increased proportions of Ki67-positive proliferating cells (Figure 7F). EdU injections were used to trace dividing cells from P0 to P1, but no significant difference in the number of cells proceeding through S phase was detected between *Peanut^−/−^* and *Peanut^+/−^*retinas, as marked by EdU after the 24-hour pulse (Figure S10). Overall, loss of *Peanut* results in altered RPC cell cycle progression, leading to increased proliferative, self-renewing divisions and decreased neurogenesis in the developing retina. Interestingly, this prolonged proliferation does not lead to any changes in cell fate or retinal morphology. Instead, the altered timing of neurogenesis may cause a developmental delay resulting in altered retinal gene expression programs during development that manifest as decreased visual function in the mature animal (Figure 5).

In summary, we show that *Peanut^−/−^* retinal RPCs have shortened cell cycle phases, biasing towards proliferative divisions, and mature retinas have decreased expression of photoreceptor genes resulting in decreased visual function. Overexpression of *Peanut* resulted in a lengthening of G2 cell cycle phase and bias towards neurogenic divisions of RPCs, as well as a promotion of photoreceptor fate. Considering these phenotypes, we suggest that *Peanut* regulates the progression of cell cycle progression and establishment of photoreceptor GRNs.

## 3. Discussion

In this study, we sought to investigate the role of lncRNAs in retinal development. We leveraged RNA-seq data from sorted populations of cells at two distinct time-points of retinal development to identify and characterize differentially expressed lncRNAs between RPCs and PMCs in the early and late developing retina. We also demonstrated that lncRNAs display cell type specific expression in the developing retina as well as in retinal ganglion cell and amacrine cell subtypes. LncRNAs are differentially expressed in retinal disease conditions (Ji et al., 2024) and are implicated in the disease etiology of age-related macular degeneration (AMD), diabetic retinopathy (DR), and more (Zhang et al., 2023; Sharma and Singh 2023; Song and Kim 2021). For example, many lncRNAs are differentially expressed in retinal ganglion cell (RGC) subtypes after axonal injury (Ayupe et al., 2021). Additionally, loss of the rod master transcription factor NRL altered the expression of a population of lncRNAs expressed in developing photoreceptors (Zelinger et al., 2017). Combined with the tissue and cell type specificity of expression, lncRNAs present as attractive biomarkers for injured or diseased tissue. However, the biological functions of these differentially expressed lncRNAs remain to be determined, and it is imperative to begin to elicit the specific roles of lncRNAs in the retina during retinal development and disease conditions.

Differentially expressed lncRNAs regulate proliferation and neurogenesis in the developing cortex (Aprea et al., 2013). Therefore, we sought to determine if differentially expressed lncRNAs could also regulate proliferation and neurogenesis of RPCs in the developing retina. Here, we focused on the previously uncharacterized lncRNA, *Gm11454,* that we named *Peanut*. We demonstrate that *Peanut* is enriched in late RPCs, where it promotes neurogenic divisions by regulating cell cycle progression, particularly the length of the G2 phase to bias the neurogenic potential of RPCs. Our results indicate that high *Peanut* in early postnatal retinal development promotes photoreceptor differentiation while loss of *Peanut* impedes appropriate expression levels of photoreceptor genes. We hypothesize that *Peanut* association with the chromatin positions the lncRNA to regulate photoreceptor GRNs, however the direct mechanisms and targets of *Peanut* remain undetermined. Nuclear-enriched lncRNAs interact with chromatin to facilitate chromatin looping, chromatin modifications, and modulate the activity of the transcriptional machinery (reviewed in Clark and Blackshaw, 2014). As we determined that *Peanut* is nuclear enriched and interacts with the chromatin, identifying regions of *Peanut-*DNA interactions will elucidate the genomic loci directly modulated by *Peanut* transcript expression. Interestingly, we did not observe expression of *Peanut* within mature photoreceptors, suggesting that deficits in photoreceptor gene expression and visual function arise from alterations to retinal development in RPCs or during early photoreceptor specification/differentiation.

Evidence from *Peanut* overexpression studies suggests that *Peanut* depletes expression of its neighboring gene, *Tox2*, and may function partially via Tox2 to increase photoreceptor fate specification at the expense of other late born retinal cell types. *Tox2* is a member of the TOX High Mobility Group Box Family and has been studied primarily in the context of the immune system and T-cell biology (Seehus et al., 2015, Xu et al., 2019, Seo et al., 2019, Collins et al., 2023). Notably, Tox2 regulates T follicular helper cell development by regulating chromatin accessibility (Xu et al., 2019). It is possible that Tox2 plays a similar role in the retina, as inhibition of *Tox2* expression via CRISPRi increased photoreceptors and decreased bipolar cells. Family member TOX3 influences neuronal development via regulation of Nestin expression (Sahu et al., 2016); however, no studies to date have determined a role for TOX2 in nervous system development. Recent genome-wide association studies have identified a single nucleotide polymorphism in the *TOX2* locus linked to Major Depressive Disorder, suggesting a functional role of the *TOX2* locus within nervous system development or homeostasis (Zeng et al., 2017).

Our results suggest Tox2 does not directly control Notch activity within RPCs. Our experiments, however, only provided a readout for presence or absence of Notch reporter activity in electroporated cells and not levels of Notch activity. Active Notch signaling in individual cells also leads to lateral inhibition in neighboring cells (Kageyama et al., 2008). It is therefore possible that high Tox2 expression increased Notch activation in individual cells and promoted lateral inhibition in neighboring cells, resulting in a failure to increase the total number of Notch-reporter positive cells. Additionally, Tox2 was shown to bind and transcriptionally regulate the Notch1 locus and reinforce Notch signaling in T follicular helper cells (Xu et al., 2019) and loss of family member TOX decreased expression of Notch target genes in innate lymphoid cells (Seehus et al., 2015). We therefore hypothesize that the loss of *Tox2* expression in Peanut-overexpressing cells directly attenuates Notch1 receptor expression (Figure 4), but *Tox2* overexpression is insufficient to increase the total number of cells exhibiting high Notch activity in the developing retina.

Notch activity regulates multiple aspects of retinal development and neurogenesis including maintenance of RPCs and promotion of Müller glia fate (Jadhav et al., 2006a; Jadhav et al., 2006b; Sahu et al., 2021; reviewed in Mills and Goldman, 2017). Additionally, Notch can regulate cell cycle progression by influencing the G1/S transition (Sarmento et al., 2005; Joshi et al., 2009; Orihara-Ono et al., 2011; Alhashem et al., 2022). In our *Peanut* overexpression model, the decrease in Notch1 expression and Notch activity likely contributes to the decrease in proliferative cells, the decrease in Müller glia, increased number of photoreceptors, and the lengthened S-phase in flow cytometry experiments, likely at the expense of a shortened G1-phase. Therefore, we propose that overexpressed *Peanut* functions to inhibit *Tox2* expression and bias Notch activity to regulate RPC neurogenesis and photoreceptor fate decisions.

In *Peanut^−/−^* retinas, there is no change in *Tox2* expression or in Notch activity, which may also explain the lack of a robust cell fate phenotype in *Peanut^−/−^* retinas. While upstream Notch regulators (e.g. *Hey1, Hey2, Ldb2*) are upregulated in *Peanut^−/−^* retinas, there is no change in Notch1 expression or Notch activity. Consistently, we see no change in cell fate or S-phase length in *Peanut^−/−^* retinas. The complete deletion of the *Peanut* locus may allow for compensatory mechanisms to mediate the loss of the lncRNA, explaining the lack of a Notch phenotype. Additionally, the 25 kb removed in the *Peanut* knockout may include cis-regulatory elements for *Tox2*, thereby resulting in an abnormal *cis-*regulatory architecture, abnormal chromatin conformation, and altered regulation of *Tox2* expression. Thus, it may be beneficial to perform an acute knockdown of *Peanut* to consider if this results in a more robust phenotype and altered *Tox2* expression. Despite this lack of a change in Notch activity, and cell fate, *Peanut* is still necessary for establishing the proper proportions of proliferative and neurogenic RPCs, as well as proper photoreceptor differentiation and function. Further characterizations of both *Peanut* and Tox2 are required to determine the mechanistic link between Peanut, Tox2, and the regulation of Notch signaling in RPCs.

## 4. Materials and Methods

### 4.1 Mouse Husbandry

All mice, including the Peanut knockout line (see below) were housed in a climate-controlled pathogen-free facility with a 12:12 light/dark cycle. All experimental procedures were approved by the Institutional Animal Care and Use Committee of Washington University in St. Louis School of Medicine. CD1 mice ordered from Charles River Laboratory were utilized for electroporation experiments. Mutant mice were maintained on a C57 Bl6/J background.

### 4.2 Generation of Peanut knockout mouse line

Guide RNAs (gRNAs) flanking the *Gm11454* locus were designed and tested for off target cutting efficiency by the Genome Engineering & Stem Cell Center at the McDonnell Genome Institute of Washington University School of Medicine. Synthetic gRNAs were ordered from Integrated DNA Technologies (Coralville, IA). Recombinant Cas9 protein was purchased from The Berkeley QB3 MacroLab (UC Berkeley). The sgRNA/Cas9 protein complexes were transfected into N2A cells and validated by target amplicon next generation sequencing, as previously described (Sentmanat et al., 2022)

The Peanut mutant mouse line was generated by electroporating gRNA and Cas9 protein using the ZEN technology as previously described (Wang et al., 2017), using C57Bl/6J donor mice, an ECM 630 Square Wave electroporation system, and omitting the placement of zygotes in acidic Tyrode’s solution. Briefly, embryos are extracted, electroporated with CRISPR-Cas9 reagents and sgRNAs (5’:GATGTGGGTACCTCTGCCGCNGG; 3’: CATCACTCAAAGCTCGATGAAGG; designed and validated by Washington University in St. Louis Genome Engineering and iPSC Center), kept in culture to the 2-cell stage, then surgically transferred into plugged pseudopregnant females. Resulting litters gave rise to genotypes in the expected ratios (Table S4).

### 4.3 Chromogenic in Situ Hybridization

Probe Synthesis: PCR with M13 primers was used to amplify probe sequence from PCRII plasmid. Antisense probes to target RNAs were generated through *in vitro* transcription using 2 µL 1X RNAPol Reaction Buffer (NEB), 2 µL 10X DIG RNA Labeling Mix (Roche), 1 µL RNase inhibitor, 4 µL PCR reaction, and 1 µL RNA Polymerase (Sp6 or T7, depending on direction probe was cloned into plasmid) (NEB). Transcription reactions were incubated at 37C for 1 hour, after which an additional 1 µL RNA Polymerase was added and incubated for an additional hour. 1 µL reaction was run on gel to confirm transcription efficiency. 2.5 µL DNase Buffer (NEB) and 1 µL DNaseI (RNase Free) (NEB) were added to the reaction and incubated at 37C for 30 minutes. Probes were either directly precipitated or hydrolyzed. RNA hydrolysis was performed through incubation in 112.5 µL Solution A (50 µM DTT, 25 µM NaHCO_3_, 40 µM Na_2_CO_3_) for 20 minutes at 60C with the reaction stopped through addition of 112.5 µL Solution B (10 µM DTT, 10 µM Glacial Acetic Acid in DEPC-treated water) and chilling 5 minutes on ice. RNA was precipitated by adding 1/10th volume 4M LiCl and 2 volumes 100% EtOH and placing on dry ice for 5 minutes. The RNA was pelleted through centrifugation for 20 minutes at 14,000 rpm at 4C. The resulting supernatant was carefully removed so as not to disturb the RNA pellet. The RNA pellet was then washed in 80% EtOH in DEPC-treated water and spin repeated. The pellet was allowed to dry, then resuspended in 100 µL 10 mM EDTA and stored at –80C. Probe sequences are indicated in Table S1.

*In situ* hybridization: Whole heads (Embryonic day (E)14 – postnatal day (P) 5) or eyes (P14) were fresh-frozen directly in OCT media (Fisher) at –80C. A cryostat was used to obtain 15 μm retinal sections, and slides were stored at –80C. RNA *in situ* hybridization was performed as previously described (Blackshaw, 2012). Briefly, slides were allowed to air dry then fixed in 4% paraformaldehyde in PBS. Slides were then washed in PBS (3×5 minutes) and incubated in 0.1M triethanolamine hydrochloride + 0.27% v/v acetic anhydride solution for 10 minutes. Slides were washed in PBS (3×5 minutes) and placed in hybridization buffer (50% formamide v/v, 5X saline-sodium citrate buffer (SSC), 5X Denhardt’s Solution (Alfa Aesar), 250 μg/ml Yeast tRNA (Roche), 500 μg/ml Sperm DNA) for 2 hours. DIG-labeled antisense RNA probes were diluted in hybridization buffer. 100µL probe solution was placed on each slide, with slides sealed using siliconized coverslips. Slides were incubated overnight at 70C in an airtight chamber humidified with 5X SSC solution. Slides are incubated in 5x SSC at 65C until the coverslips fall off the slides. Slides are washed in 0.2x SSC twice at 65C for 30 minutes each and once at room temperature for 5 minutes. Slides were then washed in Buffer 1 (0.1M Tris pH7.5, 0.15M NaCl) and incubated in antibody blocking buffer (Buffer 1 + 5% heat inactivated sheep serum (MP Biomedicals, heated at 56C 30 minutes)) for 1 hour at room temperature. Slides are placed in blocking buffer supplemented with anti-DIG-AP antibody (Roche) (1:5000) at 4C overnight. Slides were next washed in Buffer 1 (3×5 minutes) and once in Buffer 3 (0.1M Tris pH9.5, 0.1M NaCl, 0.05M MgCl2). Buffer 3 was then supplemented with 0.337 mg/mL NBT, 0.175 mg/mL BCIP, and 24 mg/100mL levamisole and added to slides for antibody detection. Reactions were stopped upon color saturation by 10-minute incubation in TE buffer.

### 4.4 MS2 Tagging and Visualization

HEK293T cells were cultured in DMEM with 10% FBS and 1x Penicillin-Streptomycin. Cells were plated in a 24-well plate on coverslips and allowed to reach confluency (∼1 day after plating). Media was changed to OptiMEM for 1 hour before transfection. Per Lipofectamine3000 (Invitrogen) protocol, 1 µL Lipofectamine 3000 was combined with 25 μL OptiMEM. 1 μL P3000 Reagent and 500 ng DNA were combined with an additional 25 µL OptiMEM. DNA included pCAGEN-Gm11454-12xMS2v6 (cloned from pSL-MS2-6X, a gift from Robert Singer (Addgene plasmid # 27118; http://n2t.net/addgene:27118; RRID:Addgene_27118) (Bertrand et al 1998) and pCAGEN-MS2 Coat Protein-GFP (pMS2-GFP was a gift from Robert Singer (Addgene plasmid # 27121; http://n2t.net/addgene:27121; RRID:Addgene_27121) (Fusco et al 2003). The two tubes were mixed and incubated for 10 minutes at room temperature. 50 µL transfection media was then added to each well of cells. After 24 hours, cells were fixed in 4% PFA for 15 minutes. Cells were washed 3×5 minutes in PBS, then incubated in DAPI 1:3000 in PBS + 0.1% Triton X-100 (PBST) for 10 minutes at room temperature, then washed again 3×5 minutes in PBST. Coverslips were then removed from each well and placed on a slide over a drop of Vectashield Hardset Mounting Media (VectorLabs). Images of MS2 slides were taken on the Zeiss LSM800 Confocal Laser Scanning Microscope.

### 4.5 Cellular Fractionation

NIH3T3 cells were cultured in DMEM with 10% FBS and 1x Penicillin-Streptomycin. Cells were plated on 12-well plates. Media was changed to OptiMEM for at least 1 hour before transfection. Cells were then transfected with pCAGEN-Gm11454 and pGFP-puro, pCAGEN and pGFP-puro, or not transfected following Lipofectamine 3000 protocol as described above. After 48 hours, the media was changed back to complete media (DMEM+FBS+Pen/Strep). After an additional 48 hours, cells were taken for fractionation: Media was aspirated, and cells were incubated in 500 μL Trypsin for 3 minutes. 500 μL complete media was added, and cells were transferred to a 15 mL falcon tube. Subcellular fractionation was performed as described in Vance et al., 2014: cells were pelleted at 400g for 10 minutes at 4C, then washed in PBS and spun for an additional 5 minutes. Cells were resuspended in 250 μL Lysis buffer (15 mM HEPES pH7.5, 10 mM KCl, 5 mM MgCl2, 0.1 mM EDTA, 0.5 mM EGTA, 250 mM Sucrose, 0.4% Igepal, 1 mM DTT, 40 U/ml RNaseOUT (Invitrogen), protease inhibitor cocktail [Roche]) and incubated 20 minutes on ice. Cells were spun at 2,000g for 10 minutes at 4C and supernatant was collected as the cytoplasmic fraction. Nuclei were resuspended in 50 μL Nuclei Lysis Buffer (10 mM HEPES pH7.5, 0.1 mM EDTA, 0.1 mM EGTA, 1 mM DTT, 40 U/ml RNaseOUT (Invitrogen), protease inhibitor cocktail [Roche]) and incubated 5 minutes on ice. Nuclei were then spun for 5 minutes at 17000g, 4C. Supernatant was collected as the nucleoplasmic fraction. Pellet was then resuspended in 50 µL Salt Extraction Buffer (25 mM HEPES pH7.5, 10% glycerol, 420 mM NaCL, 5 mM MgCl2, 0.1 mM EDTA, 1 mM DTT, 40 U/ml RNaseOUT (Invitrogen), protease inhibitor cocktail [Roche]) and incubated 30 minutes, rotating, at 4C. The sample was then spun at 17,000g for 20 minutes at 4C and supernatant was collected as the Salt Extracted fraction. The remaining pellet was resuspended in 50 μL Salt Extraction Buffer and kept as the chromatin fraction. RNA was then isolated from each fraction using the Qiagen Mini RNeasy Kit and stored at –80C. Approximately 25 ng of each RNA sample was used for cDNA synthesis following the Superscript IV kit (Invitrogen). cDNA was diluted and normalized between samples for qPCR analysis.

### 4.6 Electroporation

DNA Preparation: Phenol:chloroform:isoamyl alcohol (25:24:1) was added to 100 μg DNA solution in a 2:3 ratio, mixed through tube inversions by hand and centrifuged at 11,000g for 5 minutes. The aqueous layer was taken and DNA precipitated by adding 10% v/v 3M sodium acetate pH 5.4 and 2.5x v/v 100% ethanol. DNA was pelleted through centrifugation for 30 minutes at 15,000g and 4C. The supernatant is removed, and the pellet washed with 70% ethanol and a 5-minute centrifugation (15,000g at 4C). The ethanol is then removed, the pellet allowed to air dry for less than 5 minutes, and the DNA is resuspended to a concentration of 1 μg/μL (100μl) for *ex vivo* electroporation or 5 μg/μL (20μl) for *in vivo* electroporation. For *in vivo* electroporation, Fast Green dye is added for visualization during injection.

*Ex vivo* electroporation: *Ex vivo* electroporation was performed as previously described with minor alterations (Matsuda and Cepko 2004). Retinas were dissected from enucleated eyes in ice cold 1X PBS, leaving the lens in place. Electroporation was performed in an 8 x 5 x 3mm platinum plate electroporation chamber (Bulldog Bio, cat # CUY520P5), in 1 μg/μl DNA solution using 5 square-wave pulses (50V, 50ms pulses with 950ms intervals) from a Harvard apparatus square-wave electroporator (BTX ECM 830, Catalog No. 45-0662). Retinas are then flat-mounted on Cyclopore Track Etch-Membrane (Cytiva Whatman, 0.4 um, 13 mm diameter) with the presumptive photoreceptor layer face down on the filter. Filters are then floated in 24-well plates on 1ml of culture media (DMEM – high glucose, Sigma-Aldrich Cat. No. D5796) + 10% FBS + 1X Pen/Strep) and cultured at 37C in 5% CO_2_ for durations specified in each experiment.

*In vivo* electroporation: *In vivo* electroporation was performed as previously described (de Melo and Blackshaw, 2018). Briefly, P0 mouse pups were anesthetized on ice. The eye was opened using a sharp beveled 30-gauge needle. A 33-gauge blunt needle containing DNA solution is inserted into the subretinal space through a hole generated by a 30-gauge syringe. 0.3 µL of DNA solution is slowly injected into the subretinal space. The electroporation tweezertrode was applied to the head of the injected pup, with the positive pole electrode over the injected eye and negative pole over the noninjected eye, and electrical pulses applied (5, 90 V, 50 ms square-wave pulses with 950 ms intervals).

### 4.7 Immunostaining

Retinas were dissected fresh in ice cold PBS and fixed in 4% paraformaldehyde in PBS for 1 hour at 4C. Retinas were washed in PBS and placed in 30% sucrose in PBS. Retinas were then embedded in OCT compound (Fisher) and frozen at –80C. 15μm sections through the retina were then taken using a cryostat, and slides were stored at –20C. Prior to staining, slides were allowed to air dry for 20-30 minutes and incubated in 1X PBS for 5 minutes. Sections were then placed into Blocking Buffer (1X PBS, 5% Horse Serum, 0.2% Triton X-100, 0.02% sodium azide, 0.1% BSA) for 2 hours at room temperature. Sections were incubated overnight at 4C with primary antibody diluted 1:200 (unless otherwise noted) in Blocking Buffer. Sections were washed 3×5 minutes in 0.1% PBST and incubated 2 hours in secondary antibody 1:500 in Blocking Buffer. Sections were then incubated in DAPI 1:3000 for 15 minutes and again washed 3×5 minutes PBST. Slides were then coverslipped with VectaShield Hardset Mounting Media (VectorLabs). Images were taken on the Zeiss LSM800. Antibodies used are shown in Table S3.

### 4.8 EdU Incorporation Analysis

For EdU analyses of electroporated retinal explants, 250 μL of 50 μM EdU in complete media was added to each well to a final concentration of 10 μM 24 hours after electroporation. For P0 *in vivo* EdU analyses, mouse pups were injected subcutaneously with 20 μL of 4 mM EdU. EdU staining was performed using the Click-IT EdU AlexaFluor 647 imaging kit (Invitrogen), and slides were placed directly into primary antibody for immunohistochemistry following EdU detection protocol.

### 4.9 Flow cytometry

Retinas were dissected fresh in ice cold PBS and placed in 1 mL DMEM + 10% FBS + 1x Penicillin-Streptomycin. 250 μL of 50 μM EdU in complete media was added to final concentration of 10 μM and retinas were incubated for 2 hours at 37C. Retinas were then dissociated in 200 μL HBSS + 200 μL Papain solution (0.6mg/mL cysteine-HCL, 1mM EDTA, 0.6 mM β-mercaptoethanol, 1 mg/mL Papain) at 37C for 10 minutes, with occasional tapping. 600 μL DMEM+ 10%FBS + 1x Pen/Strep was added with 5 μL DNAseI. Cells were then incubated for 10 minutes at 37C. Single-cell suspensions were generated through gentle pipetting 10x with a P1000. Cells were then pelleted through centrifugation for 5 minutes at 500 RCF.

Supernatant was removed and cells were resuspended in 4% paraformaldehyde in PBS and incubated at room temperature for 20 minutes. Cells were spun down, supernatant removed, and cells resuspended in 100 μL 0.1% saponin, 0.5% BSA in PBS. Cells were incubated for 15 minutes at room temperature, then 500μL Click-IT Reaction cocktail (Invitrogen) was added and incubated for 30 minutes in the dark. Cells were spun down again, supernatant removed, and cells resuspended in DAPI 1:1000 in 0.1% saponin, 0.5% BSA in PBS. Cells were incubated for 15 minutes at room temperature in the dark. Cells were again spun down, resuspended in PBS, and taken for Flow Cytometry on a LSR Fortessa system. Gating was performed as shown in Figure S6. Cell cycle analysis was performed using FlowJo v10.10.0 based on DAPI staining of DNA content.

### 4.10 CD4 sorting

Retinas were dissected and dissociated in Papain solution as described above. After final centrifugation, cells were resuspended in 1 mL DMEM (no FBS or Pen/Strep). 25 μL CD4 magnetic beads (Dynabeads CD4, Invitrogen REF1145D) were washed in 1 mL DMEM, placed on magnet for 1 minute to collect the beads, and supernatant removed. Washed beads were then resuspended in 25 μL DMEM. 25 μL of beads were added to the resuspended retinal cells and rotated for 1 hour at 4C. Tubes were placed on a magnetic tube holder for 2 minutes. The CD4 (-) population was taken as the supernatant, placed back on tube to remove residual beads, then placed in the new tube. Cells were spun 1-minute 1500 RCF then resuspended in 1 mL TRI reagent (Sigma Aldrich). The CD4 (+) population of cells bound to the beads were resuspended in 1 mL DMEM and placed on magnet for an additional 2 minutes. Supernatant was removed and cells were resuspended in 1 mL TRI reagent. Cells in TRI reagent were then used for RNA Extraction.

### 4.11 RNA Extraction

Retinas or cells were placed in 1 mL of TRI reagent and homogenized. 200 μL chloroform was added; samples were mixed by vortexing for 1 minute, then incubated at room temperature for 3 minutes. Samples were centrifuged for 10 minutes at 4C at maximum speed. The aqueous phase containing RNA was then transferred to a fresh tube without disturbing the interphase. Equal volume 70% ethanol in DEPC-treated water was then added to the aqueous phase and placed on a Zymo RNA Clean and Concentrator kit column. Samples were spun 1 minute at 8000 rpm at 4C, then washed twice with 500 μL Zymo kit Wash Buffer. Samples were then washed with 70% ethanol in DEPC-treated water and spun 2 minutes at maximum speed to remove residual ethanol. RNA was then eluted in RNase-free water and stored at –80C.

### 4.12 Reverse Transcription and qRT-PCR

RNA samples were reverse transcribed following the Superscript IV Reverse Transcriptase (Invitrogen) protocol using kit reagents. cDNA was appropriately diluted for qPCR. Master mixes were prepared on ice made with 10 μL/well Sybr Green (Bulldog Bio), 1 μL/well each 5’ and 3’ primer (10 µM), and water up to 18 μL/well. 2 μL cDNA was placed in each well, followed by 18 µL master mix. Each cDNA/master mix combination was run in triplicate. The plate was sealed, vortexed, and spun down. Samples were run using a two-step 40-cycle protocol on a Bio-Rad CFX96 Thermal Cycler (Bio-Rad) or a StepOne Plus Real Time PCR system (Applied Biosystems). Primer sequences are included in Table S2. Comparison of the relative abundance of target transcript expression was normalized to *GAPDH* controls, with fold enrichment calculated based between experimental and control conditions. Statistical significance was determined using a student’s t-test on GraphPad Prism.

### 4.13 Tox2 CRISPRi design

pHR-SFFV-dCas9-BFP-KRAB was a gift from Stanley Qi & Jonathan Weissman (Addgene plasmid # 46911; http://n2t.net/addgene:46911; RRID:Addgene_46911). The dCas9-BFP-KRAB transgene was then cloned into a pCAG backbone. The *Tox2* sgRNA sequence was designed to target the end of the first exon of *Tox2 (5’ –* GCGTCTCTGCAGAAGGTAAGC– 3’), then cloned into the pmU6 plasmid (pmU6-gRNA was a gift from Charles Gersbach; Addgene plasmid # 53187; http://n2t.net/addgene:53187; RRID:Addgene_53187). A non-targeting sgRNA (5’ – GCACTACCAGAGCTAACTCACGG – 3’) was utilized as a control for *Tox2* CRISPRi experiments.

### 4.14 Electroretinogram (ERG)

ERGs were performed as previously described (Zheng et al 2023) on 2-month-old mice using the UTAS-E3000 Visual Electrodiagnostic System (LKC Technologies Inc, MD). Mice were dark-adapted overnight prior to the tests. Pupils were dilated with 1% atropine sulfate solution (Bausch and Lomb), and body temperature was kept at 37 C during the tests. Platinum 2.0 mm loop electrodes were placed on the cornea of each eye. A reference electrode was inserted under the skin of the mouse’s head, and a ground electrode was placed under the skin near the tail. Retinal responses to full-field light flashes (10 μs) of increasing intensity were recorded.

### 4.15 H&E Staining

Eyes for histological preparations were marked with a corneal tag indicating the ventral retina and fixed in 4% PFA overnight at 4C. Eyes were then washed for 5 minutes in 1X PBS and stored in 70% Ethanol for histological preparations. Eyes were paraffin embedded and sectioned using a vibratome. Central sections through the optic nerve were utilized for hematoxylin and eosin (H&E) staining.

Sections were submerged in xylenes twice for 10 minutes, 100% alcohol twice for 3 minutes, 95% alcohol twice for 3 minutes, and 70% alcohol once for 3 minutes. The well was washed in distilled water after each following staining: sections were then submerged in Hematoxylin 560 (Surgipath) for 5 to 10 minutes. After a well wash, sections were submerged for 30 seconds to 1 minute in Define working solution (½ cap Define concentrate (Surgipath) in 250 mL 75% alcohol). The well was washed once more, and sections were submerged in Blue Buffer 8 working solution (1 cap Blue Buffer 8 concentrate (Surgipath) in 250 mL distilled water) for 1 minute. After a well wash, sections were washed in 80% alcohol for 1 minute, then submerged in Eosin Y Alcoholic (Richard-Allan Scientific, REF 71204). The well was washed again, then sections were washed in 95% alcohol twice for 10 to 20 dips, 100% alcohol twice for 10 to 20 dips, then submerged in xylenes twice for 5 minutes. Sections were placed in a third xylenes, then coverslipped with Cytoseal 60 (Richard-Allan Scientific 8310-4).

Retinal sections were imaged on Zeiss Axio Observer inverted microscope coupled with an Axiocam 208 color camera (Zeiss) and retinal thickness measurements were taken 1500 um both ventrally and dorsally from the optic nerve head using Fiji software. Measurements were graphed using GraphPad Prism and significance between each data point was determined using a 2-way ANOVA analysis.

### 4.16 RNAseq library preparation and analysis

RNA was extracted from *Peanut* +/– and −/− retinas as described above. RNA library preparation was then performed following the SMARTer Stranded Total RNA High Input (RiboGone Mammalian) kit (Takara). Illumina TruSeq stranded primers were used for total RNA capture and purification. RNA Library concentrations were measured using a Qubit fluorometer and pooled for sequencing. Libraries were sequenced on shared lanes of a NovaSeq6000 for 300 cycles with 20 million targeted reads per library.

Raw sequencing reads were trimmed of adapters and low-quality bases using Trim Galore (https://www.bioinformatics.babraham.ac.uk/projects/trim_galore/). Trimmed reads were then aligned to the mm10 reference genome using STAR (Dobin et al., 2013). Results were further processed using Samtools to sort and index BAM files (Li et al., 2009). HTSeq was then used to perform gene-level quantification, and deepTools was used to visualize genome-wide read coverage (Ramírez et al., 2014, Anders et al., 2015). Differential expression analysis was then conducted using edgeR within R (Robinson et al., 2010). Gene ontology analysis was completed using the PANTHER database and analysis (Thomas et al 2022).

### 4.17 Single-cell RNA-sequencing analysis

Count matrices of previously published single-cell RNA-sequencing datasets of the developing mouse (Clark et al., 2019), mouse retinal ganglion cells (Tran et al., 2019), or mouse amacrine cells (Yan et al., 2020) were imported for downstream analyses into Monocle3 (Trapnell et al., 2014; Qiu et al., 2017a; Qiu et al., 2017b; Cao et al., 2019) from GEO using accession numbers GSE118614, GSE133382, GSE149715, respectively. Gene biotype was imported using biomaRT (Durinck et al., 2005; Durinck et al., 2009). Transcript expression for each cell was normalized for cellular read depth using transcripts per 10,000 transcripts. The number of cell types for which each transcript was expressed and average expression within unique cell (sub)types were calculated for every transcript within each dataset.

### 4.18 Data availability

Raw sequencing data and processed results from P0 and P22 bulk RNAseq are available on Gene Expression Omnibus (GEO) under the accession number GSE310182.

## Acknowledgements

The Authors would like to thank Federico Calegari for providing constructs utilized for generation of *in situ* probes for *Gm17566* and *C230034O21Rik*. The CBFRE-EGFP Notch reporter was a gift from Nicholas Gaiano (Addgene plasmid # 17705; http://n2t.net/addgene:17705; RRID:Addgene 17705).

This work was supported by the National Eye Institute of the NIH grants F32EY024201, R00EY027844, and R01EY03531 (BSC); R21EY023448 and R01EY020560 (SB); T32EY013360 (JEH), and P30EY002687 to the Department of Ophthalmology and Visual Sciences. Further support was received from an unrestricted grant to the Washington University Department of Ophthalmology and Visual Sciences and a Career Development Award (BSC) from Research to Prevent Blindness. We thank the Genome Technology Access Center at the McDonnell Genome Institute at Washington University School of Medicine for help with genomic sequencing. The Center is partially supported by NCI Cancer Center Support Grant P30CA91842 to the Siteman Cancer Center and by ICTS/CTSA Grant UL1TR002345 from the National Center for Research Resources (NCRR), a component of the National Institutes of Health (NIH), and NIH Roadmap for Medical Research. This publication is solely the responsibility of the authors and does not represent the official view of the NIH.

## Author Contributions

JEH: Investigation, Visualization, Writing – Original Draft

FS: Investigation

XZ: Investigation

SC: Resources

PAR: Software

SB: Conceptualization, Supervision

BSC: Conceptualization, Methodology, Supervision, Investigation, Visualization, Writing – Reviewing and Editing, Project administration, Funding acquisition

## Declaration of Interests

SB is a co-founder, shareholder, and scientific advisory board member of CDI Labs, LLC.

## Supplemental Files

**Supplemental Figure 1.**
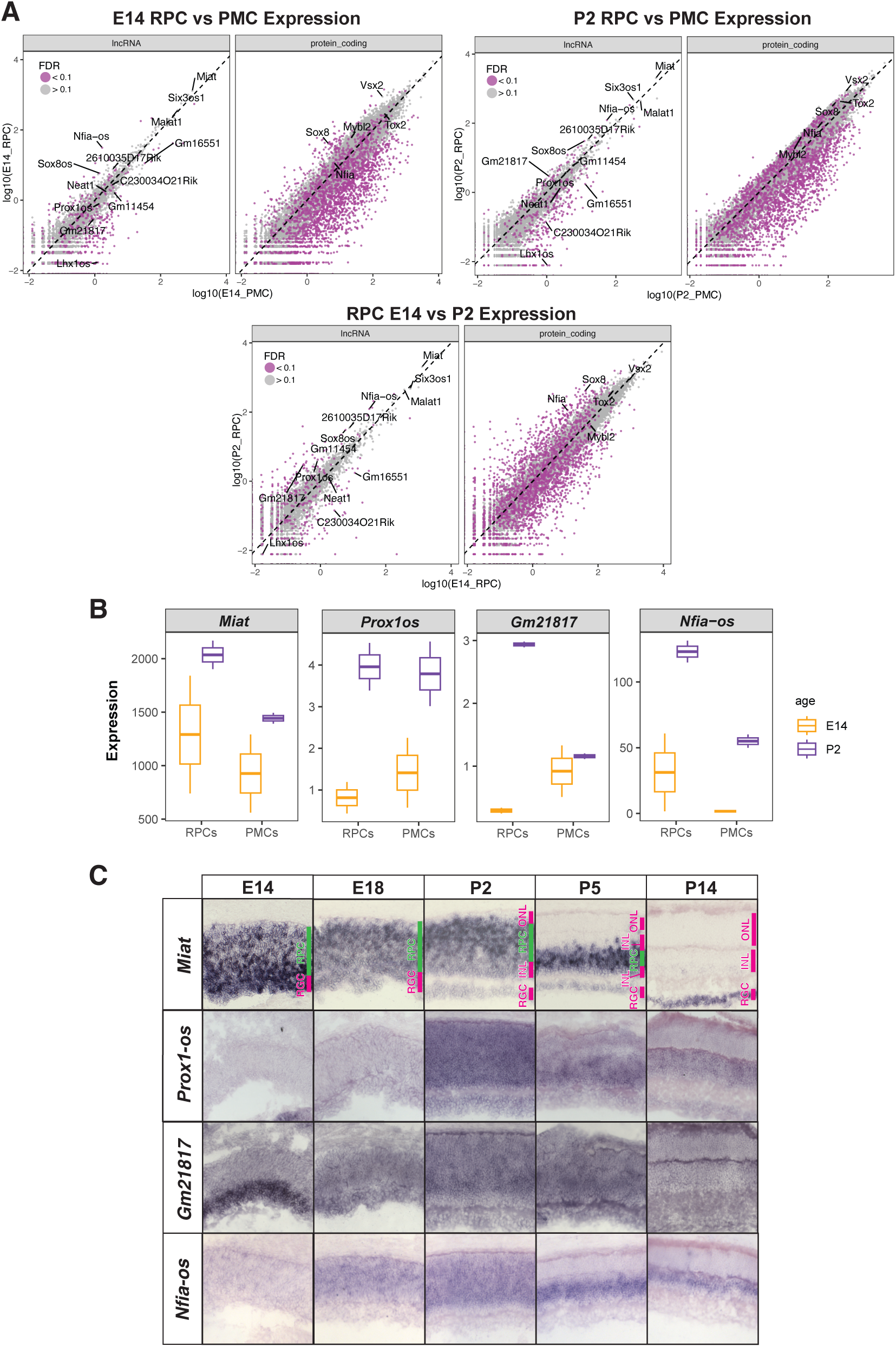
Transcript expression patterns in the developing retina. A) Differential expression of lncRNA and protein-coding transcripts across development and cell type populations (E14 RPC vs PMC, P2 RPC vs PMC, and RPC E14 vs P2). Purple color indicates differential expression across any pairwise differential expression test. B) Average expression of individual lncRNAs in RPCs and PMCs at E14 (orange) and P2 (purple). C) Chromogenic *in situ* hybridization time series confirming spatiotemporal expression patterns of individual lncRNAs.

**Supplemental Figure 2.**
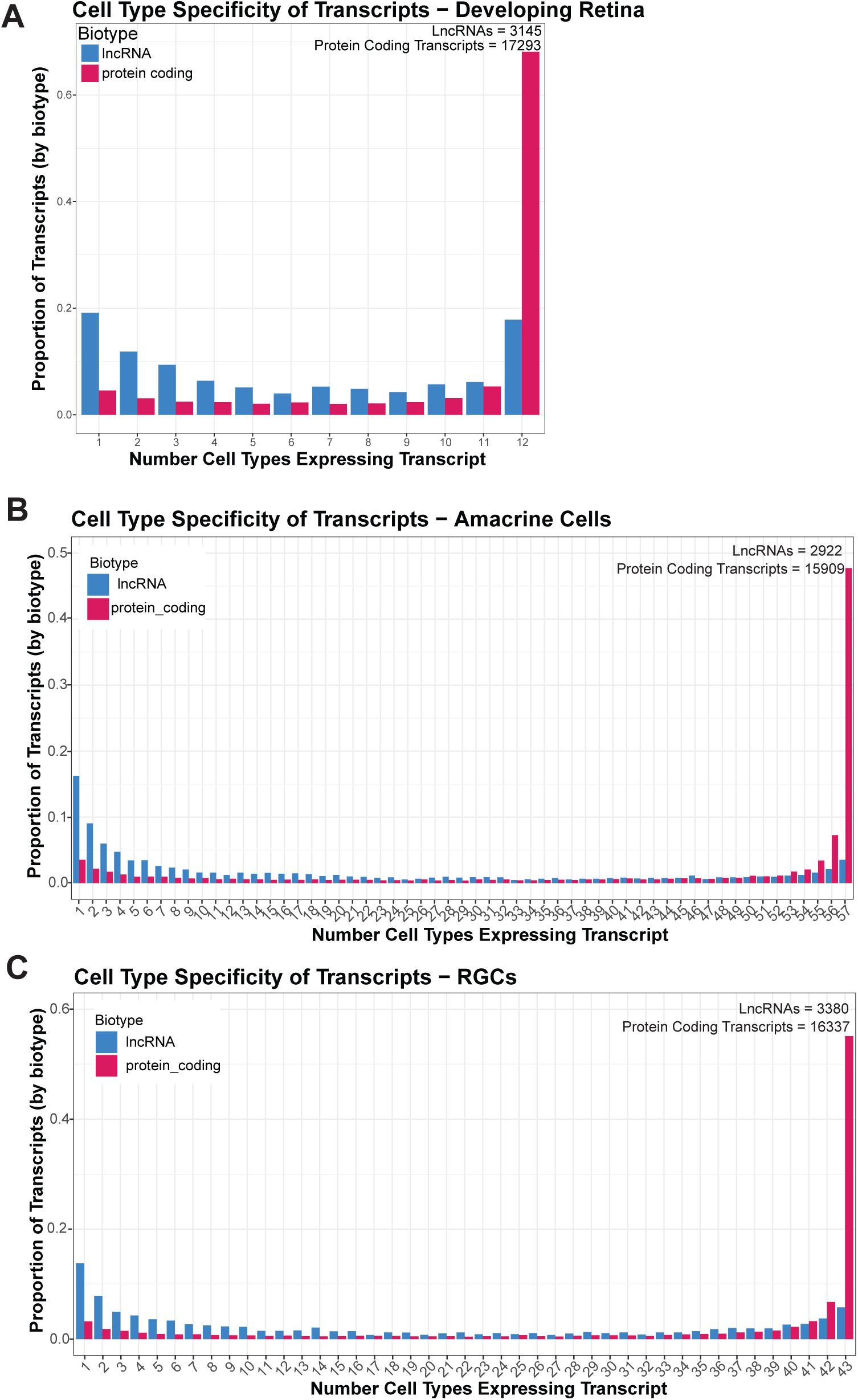
LncRNAs demonstrate more cell subtype specificity than protein coding genes. Proportion of all expressed lncRNA (blue) or protein coding (red) transcripts found in A) a given number of cell types as identified by scRNA-seq of the developing retina (Clark et al., 2019), B) a given number of retinal ganglion cell clusters (Tran et al., 2019), or C) amacrine cell clusters (Yan et al., 2020) as identified by scRNA-seq.

**Supplemental Figure 3.**
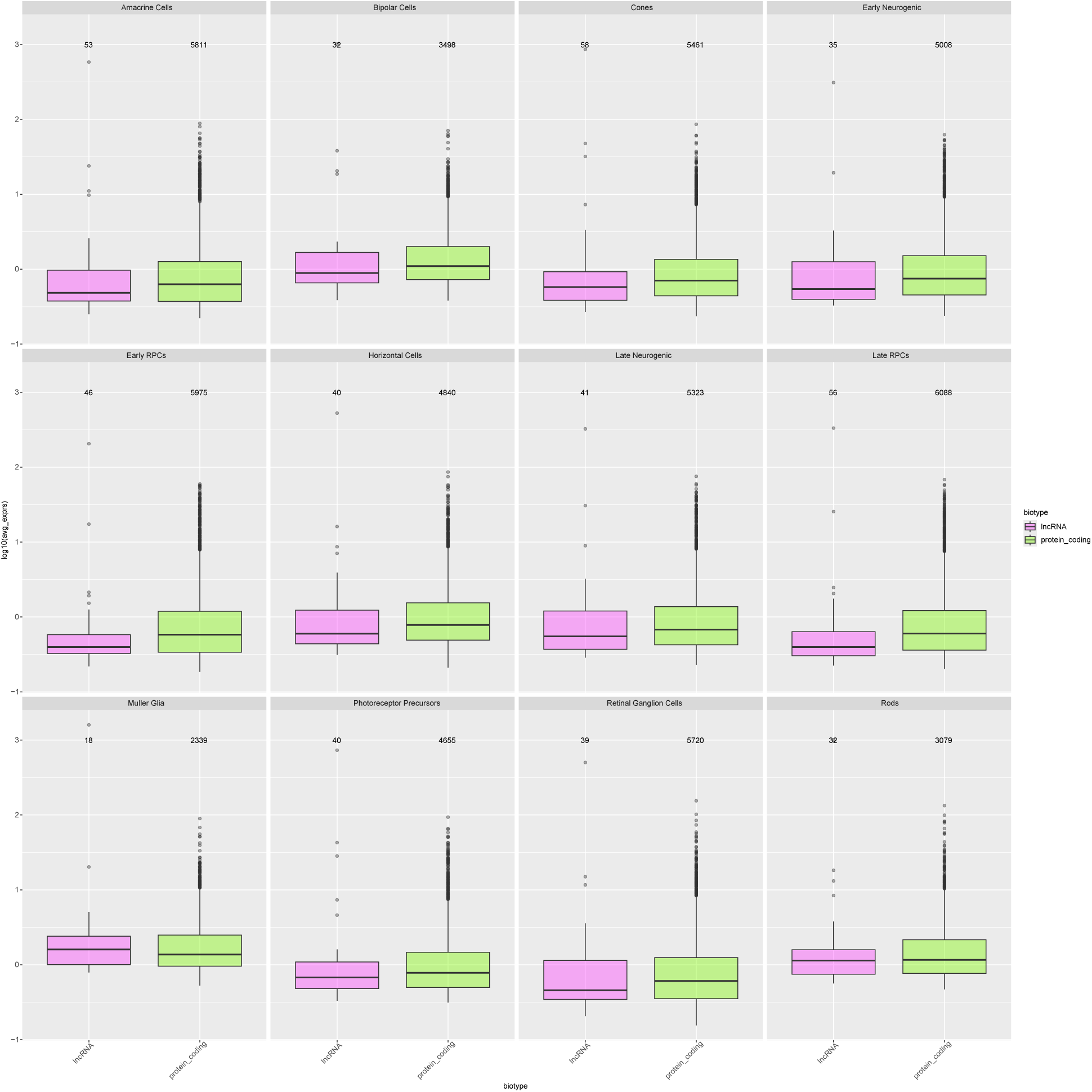
LncRNA expression levels are comparable to protein coding transcript expression levels. Average expression of lncRNA (pink) or protein coding (green) transcripts in each cell type called in previously published scRNA-seq datasets (Clark et al., 2019). Number of transcripts included shown; transcripts included were detected in at least 5% of cells for each cell type.

**Supplemental Figure 4.**
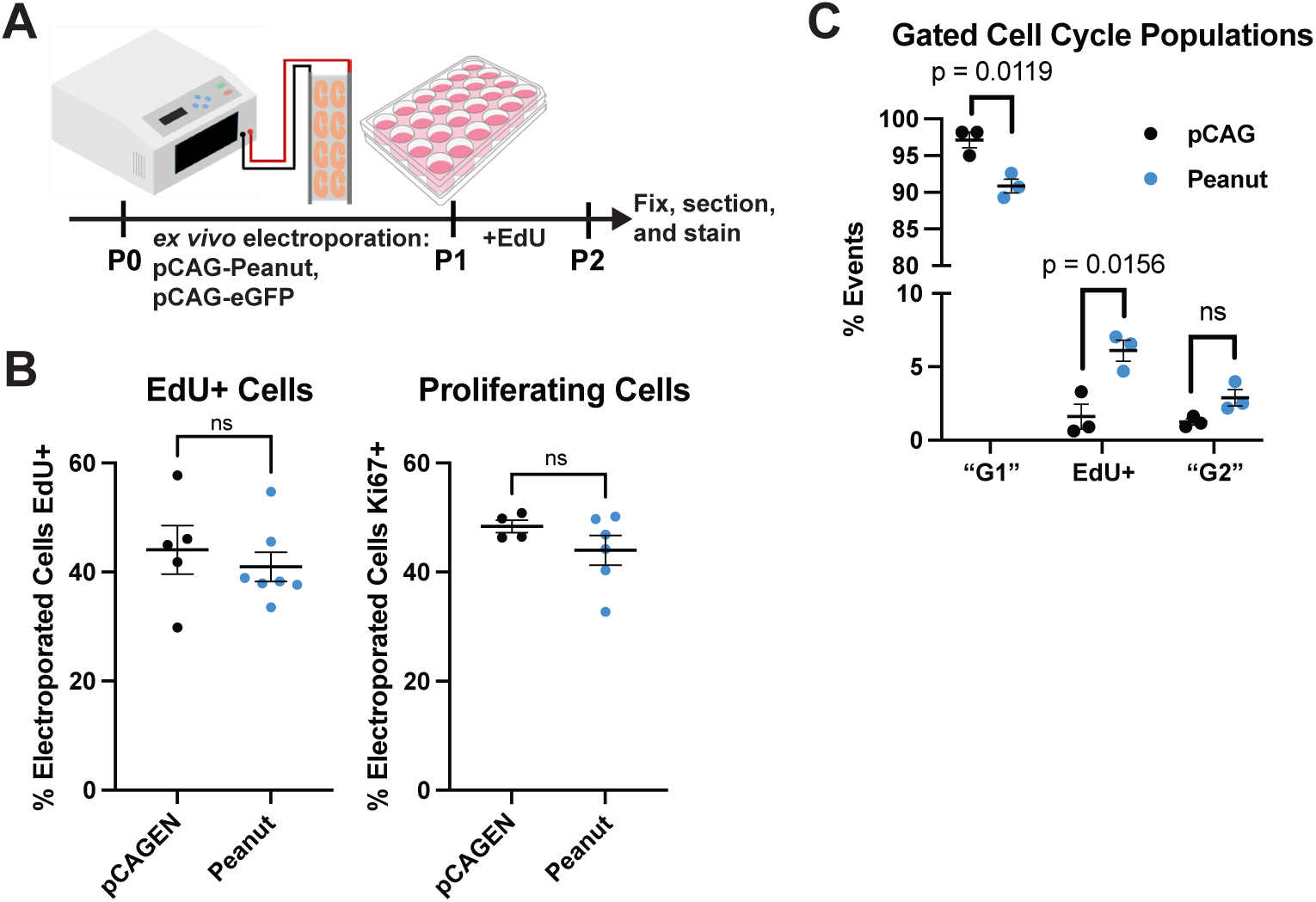
Peanut overexpression alters cell cycle progression, not cell cycle exit. A) P0 mice retinas were explanted and electroporated with pCAGEN-Peanut or empty pCAGEN and pCAG-emGFP. After 24 hours, a 24-hour EdU pulse was performed. Retinal explants were then fixed, sectioned, and stained. B) Quantification of EdU+ and Ki67+ electroporated cells after 24-hour EdU pulse. Counts were performed over triplicate images per retina. Mean ± SEM displayed, statistical significance determined by student’s t-test, ns = not significant. C) Flow cytometry analysis of DNA content in Peanut overexpression experiments indicates decreased proportions of cells in G1 and an increased proportion of cells in S phase (EdU+ after 2-hour pulse). Gating strategies shown in Supplemental Figure 5. Percent of events in flow cytometry analysis when manually gated for being EdU-, DAPI<EdU+ cells (“G1”); EdU+; and EdU-, DAPI>EdU+ cells (“G2”). Mean ± SEM displayed, statistical significance determined by student’s t-test, ns = not significant.

**Supplemental Figure 5.**
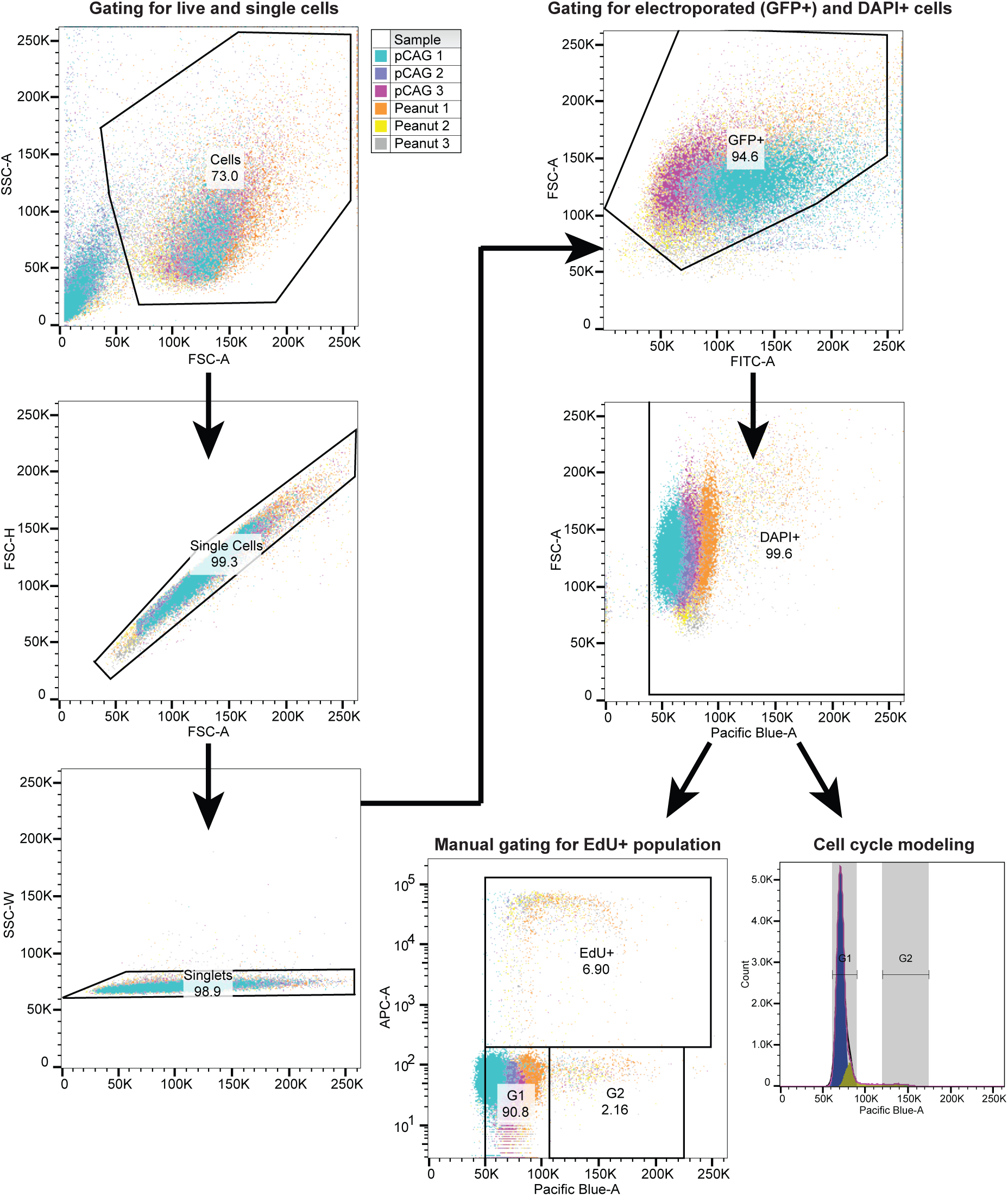
Gating strategies for flow cytometry analysis of Peanut-overexpressing retinal cells. Live cells were gated based on forward scatter area and side scatter area, then single cells were gated based on forward scatter area and height and side scatter area and width, respectively. Single cells were then gated for GFP+ (electroporated) and DAPI+ (successful staining). DAPI+ cells were then manually gated based on EdU (Fig S5) and used for cell cycle modeling in FlowJo v10 (Fig 3G). Shown are representative events from all samples with the gating and frequency for an example sample (Peanut 3).

**Supplemental Figure 6.**
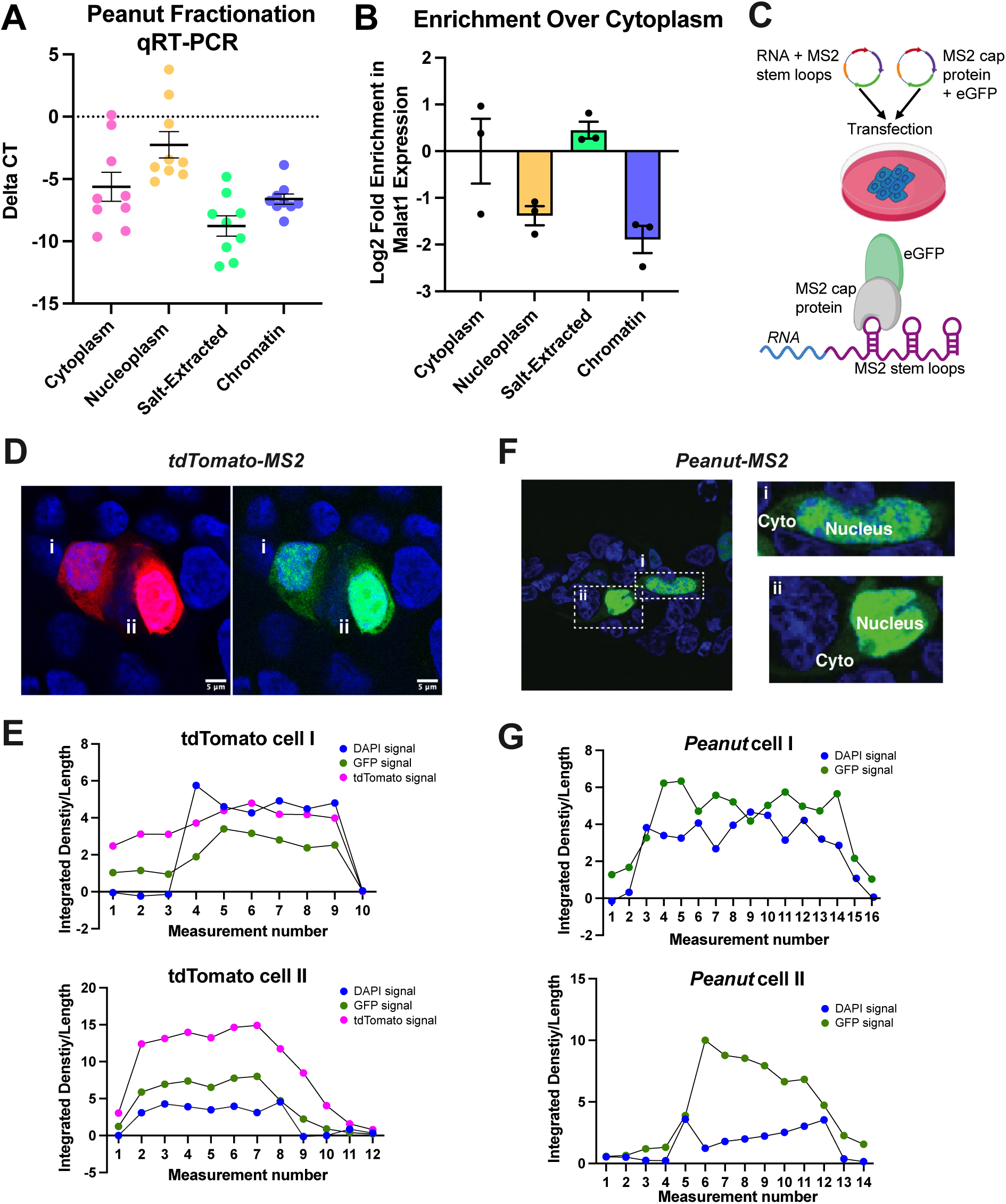
Subcellular fractionation shows enrichment of Peanut transcript in chromatin. A) Delta-CTs of *Peanut* qRT-PCR compared to GAPDH results after subcellular fractionation. B) Subcellular fractionation of NIH3T3 cells followed by qRT-PCR of *Malat1,* indicating nuclear enrichment of *Malat1* within each fraction compared to the cytoplasmic fraction. C) Control tdTomato or Peanut+MS2 stem loops and MS2 cap protein+GFP expression plasmids are transfected into HEK293 cells such that green fluorescence indicates binding of the MS2 cap protein to the MS2 stem loops of *Peanut* or *tdTomato* and red fluorescence indicates localization of tdTomato. D) tdTomato red fluorescence and MS2-GFP fluorescence showing *tdTomato* localization in HEK293 cells. E) Fiji measurements across a set length along regular intervals recording DAPI, GFP, and tdTomato fluorescence for cell i and cell ii as indicated in (D). Demonstrates GFP and tdTomato fluorescence follow similar trajectories. F) MS2-GFP fluorescence showing *Peanut* localization in HEK293 cells. G) Fiji measurements across a set length along regular intervals recording DAPI and GFP for cell i and cell ii as indicated in (F). Demonstrates the presence of some GFP signal (*Peanut* presence) in the cytoplasm (no DAPI signal), but increased signal in the nucleus as identified by increased DAPI signal.

**Supplemental Figure 7.**
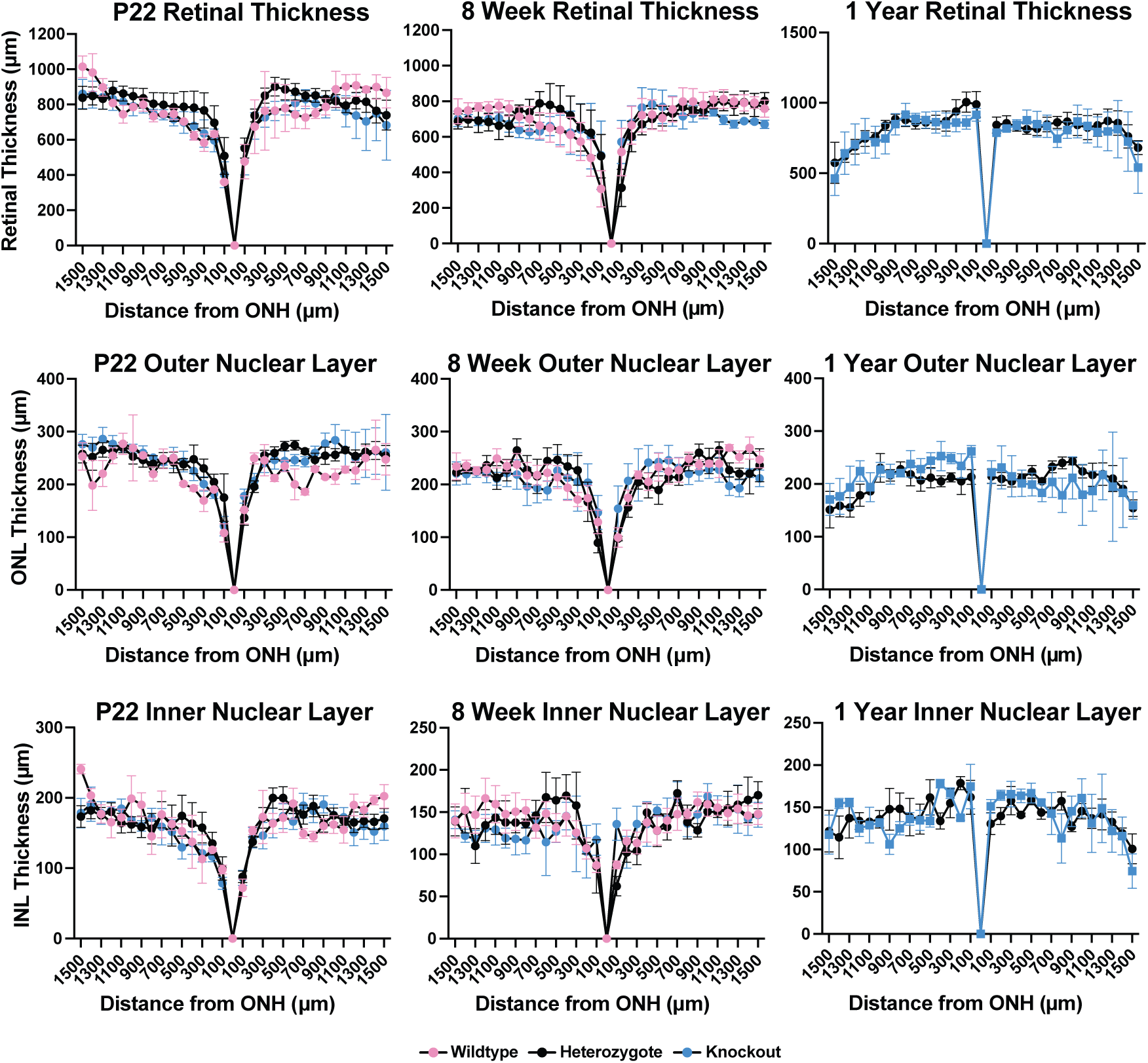
Retinal thickness of Peanut mutants. Thickness of P22, 8 weeks, and 1 year Peanut WT (Wildtype, pink), Peanut^+/−^ (Heterozygote, black), and Peanut^−/−^ (Knockout, blue) whole retinas, outer nuclear layers, and inner nuclear layers measured from H&E staining images at interval distances from optic nerve head (ONH). Shown is Mean ± SEM for each point. No significant differences were identified by 2-way ANOVA analysis.

**Supplemental Figure 8.**
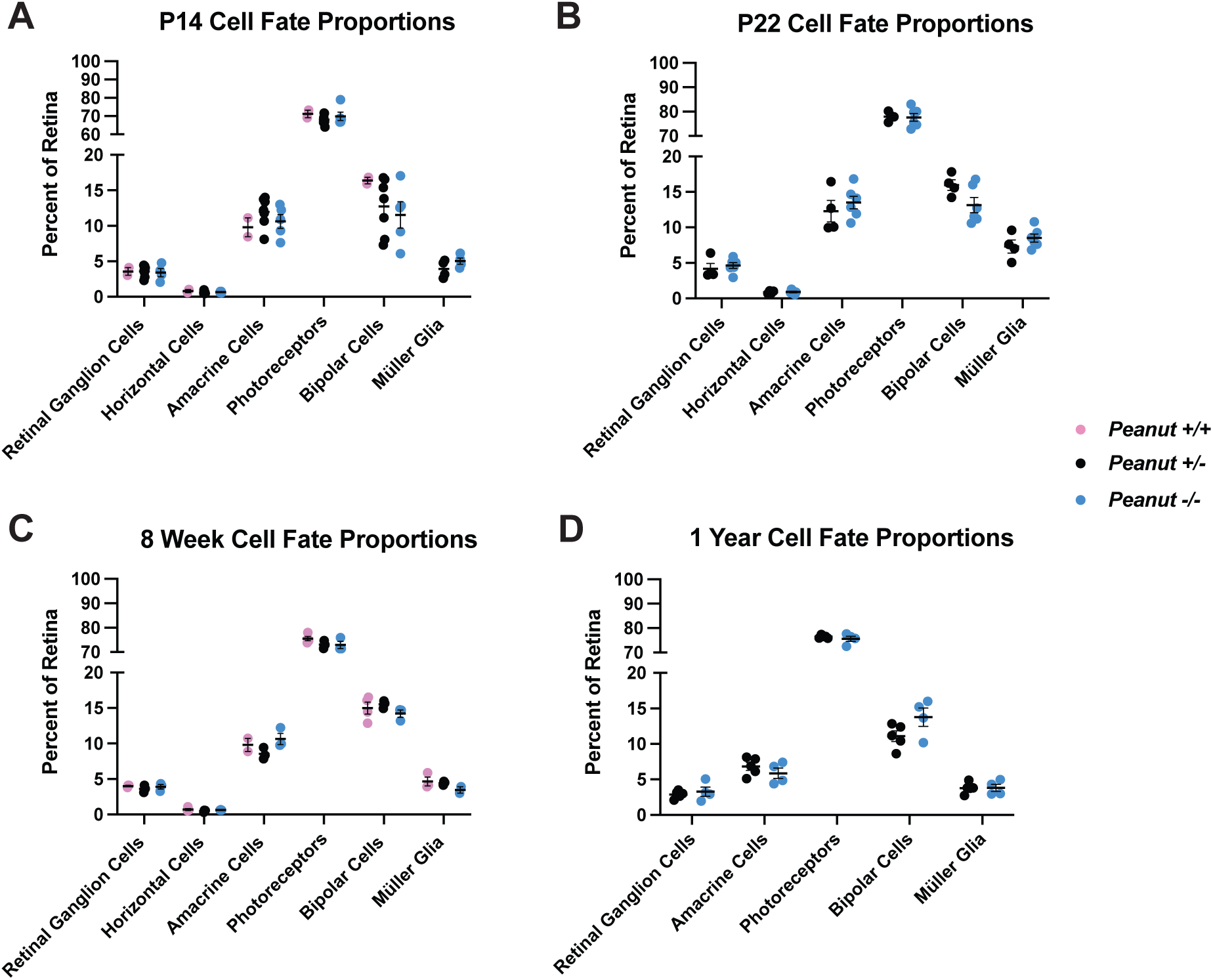
Cell fate of Peanut mutants. Mutant cell type proportions at P14, P21, 8 weeks, and 1 year. Counts were performed as percent positive cells out of total retinal cells in a set area, counts performed across triplicate images per retina. Markers used for cell populations – Retinal Ganglion Cells: RBPMS; Horizontal Cells: Calbindin; Amacrine Cells: Pax6 or Tfap2a; Photoreceptors: Crx, Otx2, or Recoverin; Bipolar Cells: Chx10/Vsx2 or Otx2; Müller Glia: p27 or Lhx2. Mean ± SEM displayed, no statistical significance was found by student’s t-test.

**Supplemental Figure 9.**
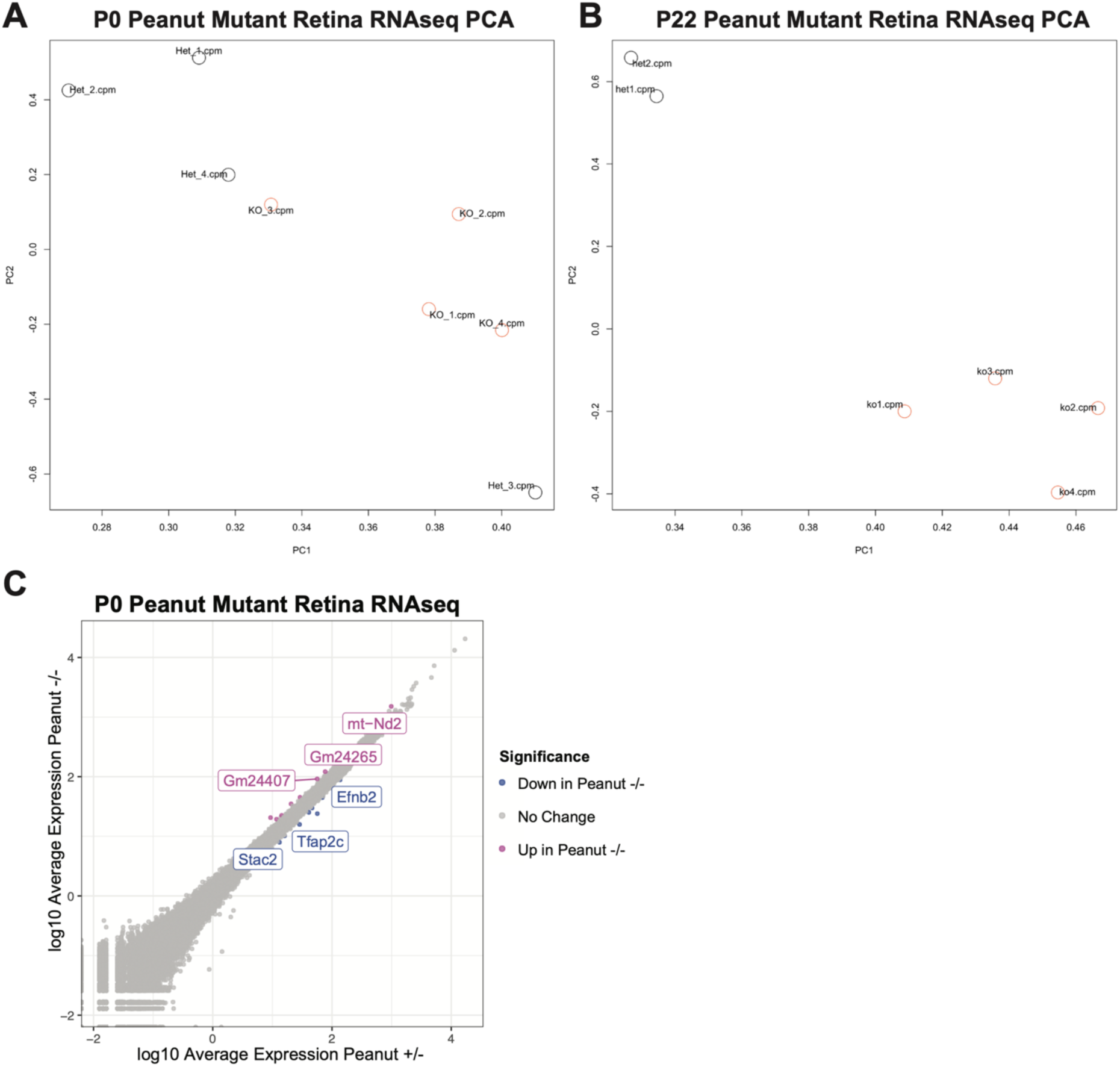
Transcript expression changes in Peanut mutants. A) Principal Component Analysis showing clustering of P0 *Peanut^−/−^ (KO) and Peanut^+/−^* (Het) retinas used for differential expression analysis. B) Principal Component Analysis showing clustering of P22 *Peanut^−/−^ (KO) and Peanut^+/−^* (Het) retinas used for differential expression analysis. C)Transcript expression in P0 *Peanut^−/−^ and Peanut^+/−^* retinas, colored by differential expression significance (FDR < 0.05) and direction (magenta=up in *Peanut^−/−^;* blue = down in *Peanut^−/−^)*.

**Supplemental Figure 10.**
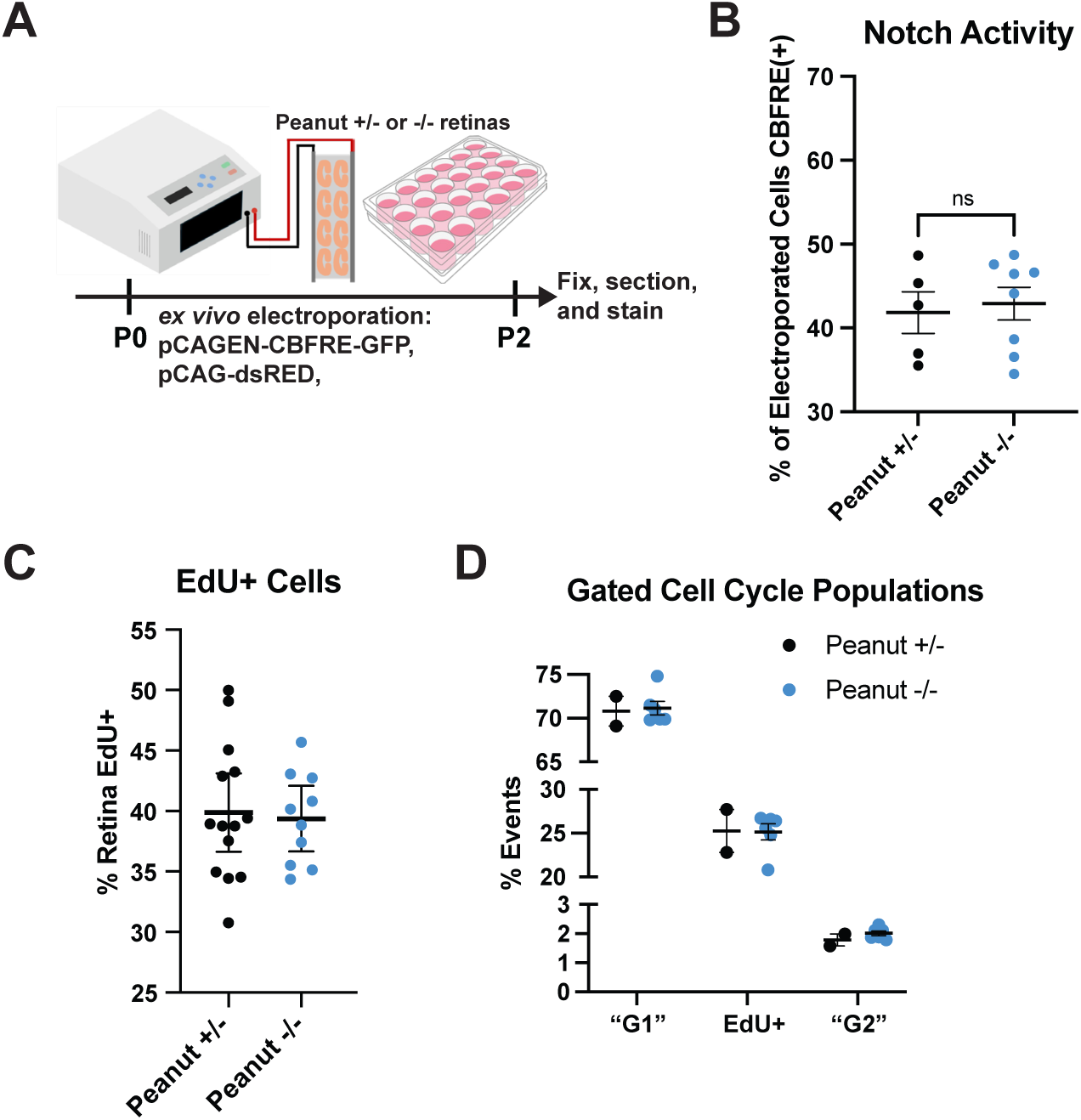
Notch signaling activity and S-phase proportions are unchanged in Peanut knockout retinas. A) P0 *Peanut^−/−^ and Peanut^+/−^* retinas were electroporated with pCAG-dsRED and pCAGEN-CBFRE-GFP, cultured for 2 days, then fixed, sectioned, and stained via immunohistochemistry. B) Quantification of electroporated cells positive for Notch Activity (CBFRE-GFP signal). Counts were performed over triplicate images per retina. C) P0 *Peanut^−/−^ and Peanut^+/−^* mice were injected with EdU for 24 hours, then retinas were taken for Click-It EdU staining. Counts were performed as percent positive cells out of total retinal cells in set area, counts performed across triplicate images per retina. D) *Peanut* knockout decreases cells in G1 and G2 but doesn’t affect cells in S phase (EdU+ after 2-hour pulse). Percent of events in flow cytometry analysis manually gated for being EdU-, DAPI<EdU+ cells (“G1”); EdU+; and EdU-, DAPI>EdU+ cells (“G2”). Gating strategy was as shown in Figure S6, with the exception of GFP+ gating as cells were not electroporated. Mean ± SEM displayed, statistical significance determined by student’s t-test, ns = not significant.

**Table S1.** In situ probe sequences. Reverse complement sequences of target genes for chromogenic *in situ* hybridization.

**Table S2.** Immunohistochemistry antibodies. Primary and secondary antibodies and dilutions used for immunostaining.

| **Antibody Name** | **Species** | **Company** | **Catalog Number** | **Dilution** |
| --- | --- | --- | --- | --- |
| Calbindin | Mouse | Sigma | C9848 | 1:200 |
| Chx10 | Sheep | Exalpha | X1180P | 1:200 |
| Crx | Mouse | Abnova | H00001406-M02 | 1:200 |
| GFP | Chicken | AVES | GFP-1010 | 1:1000 |
| GFP | Goat | Rock Island Antibodies | 600-101-215 | 1:1000 |
| GFP | Rabbit | Life Technologies | A6455 | 1:1000 |
| Ki67 | Mouse | BD Transduction Laboratories | 550609 | 1:200 |
| Lhx2 | Rabbit | Millipore Sigma | ABE1402 | 1:200 |
| Neurog2 | Mouse | Biotechne | MAB3314 | 1:200 |
| Otx2 | Goat | R&D Systems | AF1979 | 1:200 |
| p27 | Mouse | BD Transduction Laboratories | 610241 | 1:200 |
| Pax6 | Mouse | DSHB | PAX6a.a 1-233 | 1:200 |
| Phosphohistone H3 (pH3) | Rabbit | Millipore Sigma | 06-570 | 1:200 |
| RBPMs | Rabbit | GeneTex | GTX118619 | 1:200 |
| Recoverin | Rabbit | Millipore Sigma | AB5585 | 1:200 |
| Tfap2a | Mouse | Abnova | H00007070-M01 | 1:200 |
| Alexa Fluor 488 anti-chicken | Donkey | Jackson ImmunoResearch | 703-545-155 | 1:500 |
| Alexa Fluor 488 anti-mouse | Donkey | Invitrogen | A21202 | 1:500 |
| Alexa Fluor 488 anti-rabbit | Donkey | Invitrogen | A21206 | 1:500 |
| Alexa Fluor 488 anti-sheep | Donkey | Invitrogen | A11015 | 1:500 |
| Alexa Fluor 488 anti-goat | Donkey | Invitrogen | A11055 | 1:500 |
| Alexa Fluor 568 anti-rat | Goat | Invitrogen | A11077 | 1:500 |
| Alexa Fluor 568 anti-mouse | Donkey | Invitrogen | A10037 | 1:500 |
| Alexa Fluor 568 anti-rabbit | Donkey | Invitrogen | A10042 | 1:500 |
| Alexa Fluor 568 anti-sheep | Donkey | Invitrogen | A21099 | 1:500 |
| Alexa Fluor 647 anti-mouse | Donkey | Invitrogen | A31571 | 1:500 |
| Alexa Fluor 647 anti-rabbit | Donkey | Invitrogen | A31573 | 1:500 |
| Alexa Fluor 647 anti-sheep | Donkey | Invitrogen | A21448 | 1:500 |

**Table S3.**
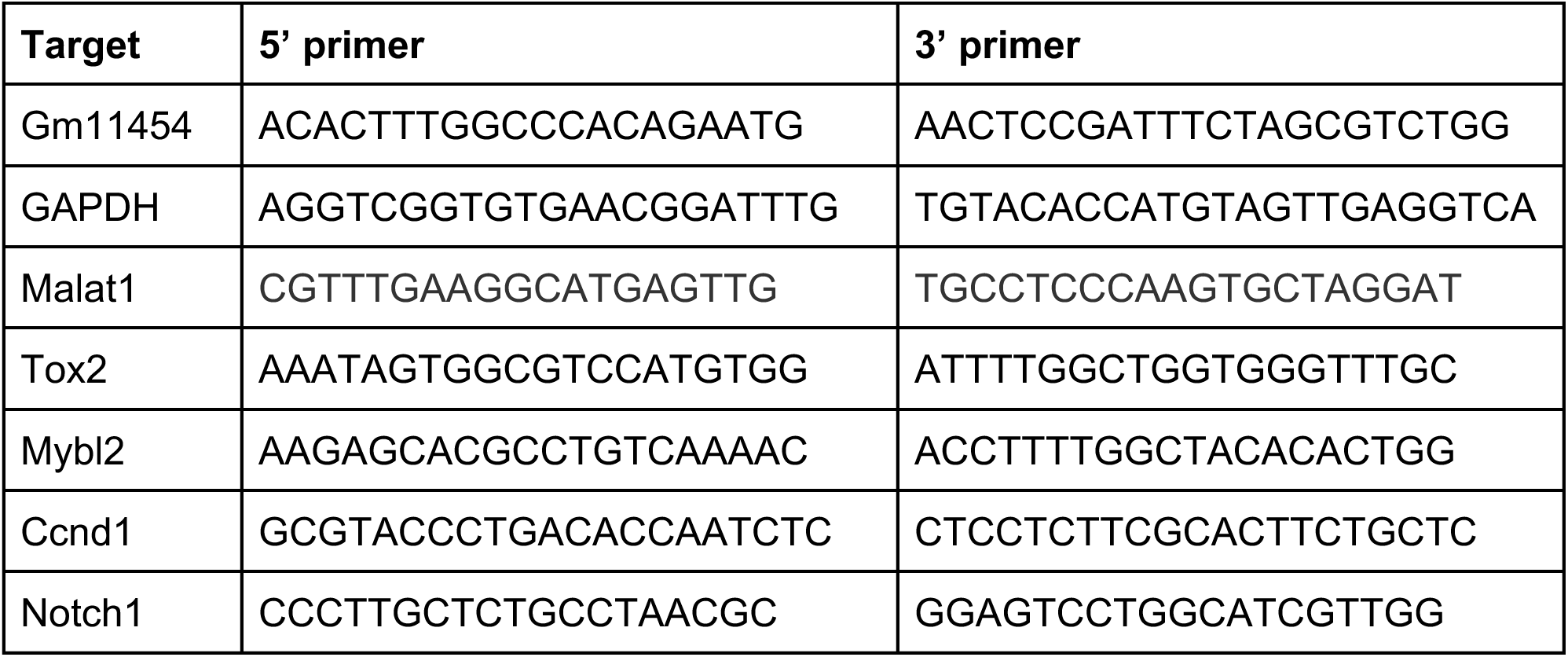
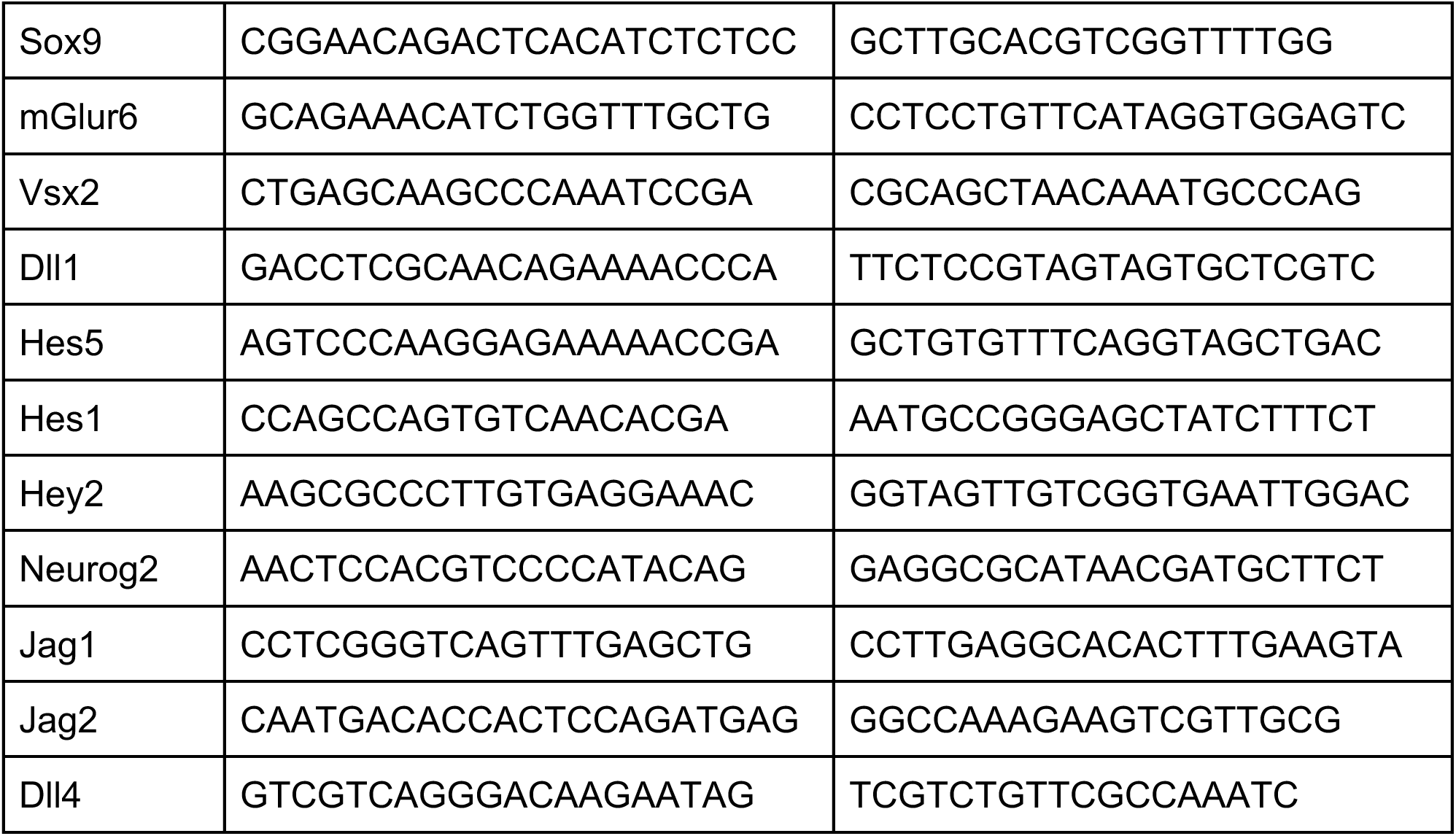
qPCR primers. 5’ and 3’ primers for target genes used in qRT-PCR experiments.

**Table S4.** Peanut mutant litter genotypes. Genotypes from heterozygote x heterozygote and heterozygote x knockout crosses were observed in expected ratios.

| Cross | Peanut+/- x Peanut+/- |  |  |  | Peanut+/- x Peanut-/- |  |  |  |
| --- | --- | --- | --- | --- | --- | --- | --- | --- |
| Genotype | Peanut WT | Peanut +/- | Peanut -/- | Unknown | Peanut WT | Peanut +/- | Peanut -/- | Unknown |
| Frequency | 0.22 | 0.43 | 0.25 | 0.10 | 0 | 0.46 | 0.47 | 0.08 |
| n (animals)= | 194 |  |  |  | 170 |  |  |  |

**Table S5.** P0 RNAseq Differential Expression Analysis. P0 *Peanut* heterozygote and knockout retinal bulk RNAseq results and EdgeR differential expression analysis.

**Table S6.** P22 RNAseq Differential Expression Analysis. P22 *Peanut* heterozygote and knockout retinal bulk RNAseq results and EdgeR differential expression analysis.

## Notes

### Competing Interest Statement

The authors have declared no competing interest.

